# The RecBC complex protects single-stranded DNA gaps during lesion bypass

**DOI:** 10.1101/2023.09.04.556180

**Authors:** Gaëlle Philippin, Pauline Dupaigne, Élodie Chrabaszcz, Maialen Iturralde, Mauro Modesti, Eric Le Cam, Vincent Pagès, Luisa Laureti

## Abstract

Following encounter with an unrepaired DNA lesion, replication is halted and can restart downstream of the lesion leading to the formation of a single-stranded DNA (ssDNA) gap. To complete replication, this ssDNA gap is filled in by one of the two lesion tolerance pathways: the error-prone Translesion Synthesis (TLS) or the error-free Homology Directed Gap Repair (HDGR). In the present work, we evidence a new role for the RecBC complex distinct from its canonical function in homologous recombination at DNA double strand breaks. We show that upon lesion encounter RecBC (independently of its catalytic activity and of the RecD subunit) is required to protect the nascent DNA in order to promote efficient lesion bypass. In the absence of RecBC, our data indicate that the nuclease ExoI can access and degrade the nascent DNA, affecting both TLS and HDGR mechanisms. We show that the recruitment of RecBC becomes particularly important at strong blocking lesions, when post-replicatively ssDNA gaps persist and are covered by the single-stranded DNA binding protein (SSB). This protective role of RecBC is reminiscent of the role of BRCA2 in protecting the nascent DNA in human cells, highlighting once again the evolutionary conservation of DNA replication mechanisms across all living organisms.

**SIGNIFICANCE STATEMENT:** In the presence of unrepaired lesions, stalled replication can restart ahead of the lesion creating a single stranded DNA gap. To preserve genome stability, this gap needs to be later filled in by one of the two lesion tolerance pathways, Translesion Synthesis or Homology Directed Gap Repair. In this study, we unveil a role for the RecBC complex in protecting the nascent DNA blocked at the lesion site from degradation by the ExoI nuclease. This protection is essential for an efficient repair of the single-stranded DNA gap. In the absence of RecBC, both lesion tolerance pathways are affected and genome stability is compromised.

## INTRODUCTION

DNA replication is constantly challenged by different types of roadblocks that can transiently slow down or stall replication progression. Unrepaired DNA lesions are one of these major roadblocks all organisms have to face. Several works in *Escherichia coli* have demonstrated that in the presence of unrepaired lesions, the progression of the replicative helicase DnaB is not halted (1–4) and replication can restart ahead of the lesion (2, 5, 6), resulting in lesion skipping and the formation of a single-stranded DNA (ssDNA) gap (7–9). In order to complete replication, the ssDNA gap is filled in by one of the two lesion tolerance pathways, identified both in prokaryotes and eukaryotes: 1) Translesion Synthesis (TLS), which employs specialized low-fidelity DNA polymerases that insert a nucleotide directly opposite to the lesion, with the potential to introduce mutations; 2) Homology Directed Gap Repair (HDGR), which uses the information of the sister chromatid to bypass the lesion in a non-mutagenic way through homologous recombination mechanisms (10–12). Regulation of these two pathways is fundamental as it defines the level of mutagenesis during lesion bypass. Our laboratory has previously shown that several factors contribute to regulate the balance between HDGR and TLS: the nature of the DNA lesion (13, 14), the amount of TLS polymerases present in the cell (15) and the density of lesions (16). Very recently, we have also shown that the processing of the ssDNA gap during the presynaptic phase of the RecF pathway plays a role in the regulation of lesion tolerance pathways (17) and that ssDNA gaps persistence at blocking lesions contributes to recruit TLS polymerases (18).

Interestingly, we have previously observed that loss of the *recB* gene has an impact on lesion tolerance pathways, as it leads to a strong decrease in TLS events, and this independently of the nuclease and the RecA loading activities of RecB (14, 19, 20). RecB is part of the RecBCD complex, also known as exonuclease V, a heterotrimeric complex that plays a role in several aspects of cell metabolism and genome maintenance, such as DNA double-strand break repair, conjugation, chromosomal segregation, degradation of foreign DNA and completion of DNA replication (21–23). In particular, RecBCD is the key factor in homologous recombination pathway at double stranded DNA breaks, also referred to as the RecBCD pathway (24–27). The complex is composed of three subunits: RecB is the main subunit, containing a DNA-dependent ATPase activity, a 3’->5’ DNA helicase activity, the nuclease active site and the RecA interacting domain (28). RecC itself does not possess any catalytic activities, but it does bind ssDNA, and can stimulate the ATPase and helicase activities of the RecB protein, with which it forms a complex of 1:1 stoichiometry (29). The main role of RecC is to recognize a specific DNA sequence, the χ (Chi) site (30–32), that participates in the regulation of the complex. RecD is the smallest subunit and possesses a ssDNA-dependent ATPase activity and 5’->3’ DNA helicase activity (33, 34). The other functions of RecD are to regulate the nuclease activity of RecB, to increase the processivity of the complex and to negatively regulate RecA loading (35–38). The preferred substrate of the RecBCD complex is a blunt or nearly blunt double stranded DNA ends: the RecB subunit binds to the 3’ end while the RecD subunit associates with the 5’ extremity (39–41). Individually, each subunit is a poor ATPase, helicase or nuclease, but together they form one of the most processive helicase/nuclease machine (41–43).

Initially, the complex was thought to contain only RecB and RecC subunits (44), and RecD was identified later as the subunit regulating the nuclease activity and containing a helicase activity (45–47). Whereas *recD* null mutants display normal cell viability, resistance to UV light and other DNA damaging agents, and are recombinant proficient, mutations in *recB* or *recC* genes strongly affect cell viability, reduce conjugal recombination 100- to 1000-fold, and sensitize cells to DNA-damaging agents (21). This mild phenotype of the recD mutant is due to the fact that in the *recD* deficient strain, the nuclease activity can be substituted by the nuclease RecJ (47) and that RecA is constitutively loaded (36). Since the single *recB* and *recC* mutant strains resulted in identical phenotypes, this suggested that the minimal complex was RecBC. However, *in vitro* RecBC and RecBCD complex do not possess the same enzymatic properties: several works showed that compared to the potent and processive nuclease/helicase RecBCD, RecBC has no exonucleolytic activity on a linear substrate and its endonucleolytic and helicase activities are strongly inhibited (28, 33, 48). Therefore, it is possible that the two complexes coexist in the cell with different physiological functions, but up to now, no specific cellular role has been associated to the sole RecBC complex.

In this work, we show that the RecBC complex is required to protect the nascent DNA when a persistent ssDNA gap is formed after lesion skipping and repriming. We propose that the RecBC complex, by binding to the SSB-covered ssDNA gap, disorganizes the ssDNA region in a way that will protect the nascent DNA from the action of the nuclease ExoI. In the absence of RecBC, ExoI action on the nascent DNA turns out to strongly reduce TLS and also to affect HGDR mechanisms. Therefore, the RecBC complex is necessary to enable efficient lesion bypass, a new function independent of its catalytic activities and its canonical role in DNA double strand break repair.

## MATERIAL AND METHODS

### Bacterial strains and growth conditions

All *E. coli* strains used in this work are listed in Table S1. They are derivatives of strains FBG151 and FBG152 (13), that carry a plasmid that allows the expression of the *int–xis* genes after IPTG induction. Strains were grown on solid and liquid Lysogeny Broth (LB) medium. Gene disruptions were achieved by the one-step PCR method (49) or by P1 transduction using the Keio collection (50). The antibiotic resistance cassette was removed according to (49) in order to avoid polar effect on the other operon genes and the entire open reading frame of the gene is removed. Following the site-specific recombination reaction, the lesion is located either in the lagging strand (FBG151 derived strains) or in the leading strand (FBG152 derived strains). The mutant alleles of *exoI* (*exoI^A183V^*) has been obtained by CRISPR-Cas9 technology adapted for *E. coli* according to (51). This strain has been verified by sequencing (the entire modified gene). Antibiotics were used at the following concentrations: ampicillin 50 or 100 μg/ml; tetracycline 10 μg/ml, kanamycin 100 μg/ml, chloramphenicol 30 μg/ml. When necessary, IPTG and X-Gal were added to the medium at 0.2 mM and 80 μg/ml, respectively.

### Plasmids

pVP135 expresses the integrase and excisionase (*int–xis*) genes from phage lambda under the control of a *trc* promoter that has been weakened by mutations in the -35 and the -10 region. Transcription from *Ptrc* is regulated by the *lac* repressor, supplied by a copy of *lacI^q^* on the plasmid. The vector has been modified as previously described (13).

pVP146 is derived from pACYC184 plasmid where the chloramphenicol resistance gene has been deleted. This vector, which carries only the tetracycline resistance gene, serves as an internal control for transformation efficiency.

pVP141-144 and pGP9 are derived from pLDR9-attL-lacZ as described in (13). pLL1 and pLL7 are derived from pVP141 as previously described (14). pAP1, pAP2 and pGP22 are derived from pVP143. All these plasmid vectors contain the following characteristics: the ampicillin resistance gene, the R6K replication origin that allows plasmid replication only if the recipient strain carries the *pir* gene, and the 5’ end of the *lacZ* gene in fusion with the *attL* site-specific recombination site of phage lambda. The P’3 site of *att*L has been mutated (AATCATTAT to AAT<u>TA</u>TTAT) to avoid the excision of the plasmid once integrated. These plasmids are produced in strain EC100D pir-116 (from Epicentre Biotechnologies, cat# EC6P0950H) in which the pir-116 allele supports higher copy number of R6K origin plasmids. Vectors carrying a single lesion for integration were constructed as previously described following the gap-duplex method (13). A 15-mer oligonucleotide 5’-ATCACCGGC<u>G</u>CCACA-3’ containing or not a single G-AAF adduct (underlined) in the *Nar*I site was inserted into the gapped-duplex pLL1/7 (to measure HDGR) or into the gapped-duplexes pVP141-142 or pVP143-144 to score respectively for TLS0 Pol V-dependent and for TLS-2 Pol II-dependent. A 13-mer oligonucleotide, 5ʹ-GAAGACCT<u>G</u>CAGG, containing no lesion or a dG-BaP(-) lesion (underlined) was inserted into the gapped-duplex pVP143/pGP9 to measure Pol IV TLS events. A 14-mer oligonucleotide 5’-ATACCCGG<u>G</u>ACATC-3’ containing no lesion or a single G-AF or a 1hpG adduct (underlined) was inserted into the gapped-duplex pAP1-AP2 to measure TLS like events. A 20-mer oligonucleotide 5’-CTACCT<u>G</u>TGGACGGCTGCGA-3’ containing or not the N^2^-furfuryl adduct was inserted into the gapped duplex pAP1-pGP22 to measure TLS like events. All the new batches of plasmid constructions are carefully quantified and validated by integration in the parental strain.

pVP148 plasmid expresses both *polB* and *umuD’C* loci under their own promoters. It is derived from pRW134 (52), a pGB2 vector that carries *umuD’C*, in which the spectinomycin resistance gene was replaced by the chloramphenicol resistance gene. The *polB* gene was added by subcloning a SalI-BamHI fragment from pYG787 (53). UmuD’ is the cleaved version of UmuD that forms the heterotrimeric complex (UmuD’)_2_UmuC, *i.e.*, the active form of Pol V (54). pLL70 is a plasmid expressing *exoI* gene under its own promoter. It was obtained by removing the *umuDC* gene and its promoter from pVP145 by PstI/BamHI digestion and cloning *exoI* gene+promoter by PstI/BamHI. pVP145 is derived from pRW134 (52), in which the spectinomycin resistance gene was replaced by the chloramphenicol resistance gene.

### Integration protocol: monitoring HDGR and TLS events

The protocol for lesion integration assay is described in details in (20, 55). Cells were diluted and plated before the first cell division using the automatic serial diluter and plater EasySpiral Dilute (Interscience) and were counted using the Scan 1200 automatic colony counter (Interscience).

For every integration experiment, we used the lesion versus non-lesion plasmid constructions, each plasmid solution containing an equal amount of pVP146 plasmid as internal control. Following the integration of the pLL1/7 vector (AAF lesion), sectored blue/white colonies represent HDGR events. Following integration of the vectors pVP141/142 (TLS0 for G-AAF), pVP143/144 (TLS-2 for G-AAF), pVP143/pGP9 (BaP lesion), pAP1-AP2 (G-AF or 1hpG lesions), pAP1-pGP22 (N^2^-furfuryl lesion) sectored blue/white colonies represent TLS events. The relative integration efficiencies of lesion-carrying vectors compared with their lesion-free homologues, normalized by the transformation efficiency of pVP146 plasmid in the same electroporation experiment, allow the overall rate of lesion tolerance to be measured (which corresponds to the cell survival in the presence of a single lesion). In the parental strain one lesion gives about 100% cell survival. For the G-AAF lesion we combine the results obtained with the different plasmids to measure HDGR and TLS events. Briefly, the mean of the absolute values (blue colonies/total colonies) of HDGR and TLS events were multiplied by the mean of the cell survival obtained with all the plasmids construction for a given lesion. Indeed, whether TLS or HDGR was measured, cell survival will not change for a given lesion. The value of DCL was calculated by subtracting the HDGR and TLS values to the total cell survival.

Tolerance events (Y axis) represent the percentage of cells able to survive in presence of the integrated lesion compared to the lesion-free control. The data in every graph represent the average and standard deviation of at least three independent experiments of a lesion inserted in the leading (or in the lagging) orientation. Statistical analyses were done using GraphPad Prism applying an unpaired *t*-test.

### RecBC protein expression and purification

The protocol for RecBC expression and purification has been adapted from (38). Briefly, *E. coli* RecBC complex was purified from BL21 DE3 Δ*recBD* strain harboring B13 (pETduet-His_6_-TEVsite-recB, Amp^R^) and B3 (pRSFduet-recC, Kan^R^) plasmids. The cells were induced with 1 mM IPTG at mid-log phase (0.6 OD_600_) and grown for 4h at 37°C. The lysate was purified on HisTrap HP Ni-Sepharose (Cytiva), then dialyzed overnight with TEV protease to cleave the His-tag. The His-tag was removed by a second step on HisTrap HP column. The complex was further purified using HiTrap Heparin-Sepharose (Cytiva), then HiTrap Q FF Sepharose (Cytiva) before continuing with MonoQ (Cytiva) anion exchange chromatography. The protein was dialyzed overnight in 20 mM Tris-HCl pH8, 1 mM EDTA pH 8, 1 mM DTT, 50 mM NaCl, 10% glycerol and flash-frozen.

### Preparation of the gapped DNA substrates

#### Plasmid with a 5.3 kb ssDNA gap

3 pmol of a 50 nt oligonucleotide (VP683: TCGAGCTGCGCAAGGATAGGTCGAATTTTCTCATTTTCCGCCAGCAGTCC) was 5’ radiolabelled using 3.3 pmol of γ-^32^P-ATP (Perkin Elmer) and 10 units of PNK (NEB) in a total volume of 25 µl PNK buffer 1X. The reaction was incubated during 30 minutes at 37°C and inactivated during 20 minutes at 65°C. Then it was purified using a MicroSpin G-25 column (Cytiva). The oligonucleotides (60 nM) were hybridized to χπX174 ssDNA plasmids (5.3 kb, NEB) (20 nM) in 50 μl buffer containing 10 mM Tris-HCl pH8, 1 mM EDTA pH8 and 50 mM NaCl. The reaction was incubated in water at 75°C and let cool down to room temperature. Substrates were checked on 0.8% agarose gel.

#### Plasmid with a 2.5 kb ssDNA gap

10 μM oligonucleotides (VP1394: ACGATTAACCCTGATACCAA) were phosphorylated with 1 mM ATP (Sigma) and 10 units PNK (NEB) in 50 μl PNK buffer 1X (30’ at 37°C, inactivation 20’ at 65°C). A 2.8 kb dsDNA fragment was obtained by PCR with Q5 Polymerase (NEB) using phosphorylated VP1394, VP1361 (ACGCCCTGCATACGAAAAGA) and χπX174 as template. PCR products were digested by ExoI (NEB) to remove excess of oligos and purified using Qiagen PCR purification kit. The phosphorylated DNA strand was then specifically digested with lambda exonuclease (NEB); the ssDNA strands obtained were checked on 0.8% agarose gel. They were 5’-radiolabelled using γ-^32^P-ATP and purified using a MicroSpin G-25 column. Then they were hybridized to χπX174 ssDNA plasmids (ratio 1 :1) in 50 μl buffer containing 10 mM Tris-HCl pH8 and 50 mM NaCl. The reaction was incubated in water at 80°C and let cool down to room temperature. Substrates were checked on 0,8% agarose gel.

This substrate (non radio-labelled) was also used for TEM analysis.

### Electrophoretic mobility shift assays

The RecBC complex was freshly expressed and purified several times since the RecBC complex proved to have a reduced shelf life as also mentioned by (48). The purity of the protein prep was verified by mass spectrometry. The SSB protein was provided by BioAcademia in his buffer (20 mM Tris-HCl 7.6, 200 mM NaCl, 1 mM DTT, 1 mM EDTA, 50% glycerol), and the truncated SSB ΔC8 protein was kindly provided by Michael Cox (University of Wisconsin-Madison). To note, we experimentally observed that the concentration of SSB required to induce a mobility shift was not proportional to the length of the ssDNA to be covered. Indeed, in the presence of 2.5 kb gapped substrate a higher concentration of SSB was required compared to the 5.3 kb gapped substrate (compare Fig. S5 and Fig. 4).

For the experiment in Fig 4B, 0.4 nM of the radiolabelled 5.3 kb ssDNA gap substrate (∼ 2 µM nucleotides of ssDNA) were incubated for 20’ at RT with increasing concentrations of RecBC (0 – 16 – 32 – 64 – 96 nM) in 10 µl binding buffer (14 mM Tris-HCl pH8, 40 mM NaCl, 100 mg/ml BSA-Ac, 0,6 mM DTT). The samples were loaded on 0.8% TAE agarose gel. The gel was dried for 1h30 at 60°C, exposed overnight to a Fuji screen and scanned with a Typhoon (Cytiva).

For the experiment in Fig 4C, 0.27 nM of the radiolabelled 5.3 kb ssDNA gap substrate (1.4 µM nucleotides of ssDNA) were first incubated for 20’ at RT with three fixed concentrations of SSB tetramer (2.2 – 4.3 – 8.7 nM) in 15 µl binding buffer (20 mM Tris-HCl pH8, 50 mM NaCl, 100 mg/ml BSA-Ac, 1 mM DTT and 1 mM Mg-Ac). Then increasing concentrations of RecBC (0 – 11 – 21 – 43 – 85 nM) were added to the reactions and incubated for 20’ at RT. The samples were loaded on 0.8% TA agarose gel, running at 4°C. The gel was dried for 1h30 at 60°C, exposed overnight to a Fuji screen and scanned with a Typhoon (Cytiva). Similar reaction conditions were used in Fig 4D except for substrate concentration (0.4 nM), SSB and SSB ΔC8 tetramer concentration (19.5 nM) and RecBC concentration (160 nM).

### Transmission Electron Microscopy (TEM)

The DNA substrate containing a 2.5 kb ssDNA gap (3.2 nM molecules, 10 μM nucleotides of ssDNA) was first incubated with two different concentrations of SSB protein (130 nM SSB for the ’lower’ concentration, 286 nM for the ’higher’ concentration) during 10’ at 37°C in a binding buffer containing 10 mM Tris-HCl pH 7.5, 50 mM NaCl, 2 mM MgCl_2_, 1 mM DTT and in presence or absence of AMP-PNP cofactor. Then increasing concentrations of RecBC protein (200, 250 and 300 nM) were added to the reaction during 10’ at 37°C. The effect of 300 nM RecBC was also tested on the same substrate (same concentration, same buffer) in absence of SSB protein. In parallel, the same reaction was carried out on a single-stranded substrate: 10 μM nucleotides of ssDNA (the χπX174 virion) was incubated with 286 nM of SSB protein during 10’ at 37°C in the same binding buffer, then 300 nM of RecBC were added to the reaction during 10’ at 37°C.

Complexes formed with or without RecBC were quickly diluted 40 times in the binding buffer then spread onto the TEM grids as following: 5 μL were immediately deposited for 1 minute on a 600-mesh copper grid coated with a thin carbon film, preactivated by glow-discharge in the presence of amylamine (Sigma-Aldrich) in a homemade device as previously described (56). The grids were washed with 0.2% (w/vol) aqueous uranyl acetate (Merck) and dried with ashless filter paper (VWR). The grids were directly subjected to TEM analysis in filtered annular dark-field mode with a Zeiss 902 microscope using Veletta high-resolution CCD camera and iTEM software (Olympus, Soft Imaging Solutions). TEM images were acquired at a magnification of x 85000. For each condition, at least 200 molecules were analyzed, 50 of which were measured.

## RESULTS

We have previously shown that in the presence of a UV photoproduct lesion (*i.e.* TT6-4), deletion of *recB* gene affects lesion tolerance pathways, decreasing HDGR mechanisms and strongly inhibiting the TLS pathway (14, 20). We have also shown that this role in lesion bypass is independent of the nuclease and the RecA loading activities of RecB and more interestingly, that the RecB contribution to lesion bypass does not involve processing of a DNA double strand break that could potentially arise in the presence of a ssDNA gap (20). In the present study, we further investigate and characterize this unexpected role of RecB.

To monitor lesion tolerance mechanisms, we took advantage of our previously described genetic assay (see Fig. S1A) that allows us to insert a single replication-blocking lesion into the chromosome of *E. coli,* either on the lagging or in the leading strand orientation regarding the replication fork direction. TLS or HDGR events are visualized and measured by a colorimetric assay based on the *lacZ* reporter gene using specific lesion integration plasmids (Fig. S1B and 1C) (13, 14, 55). This assay had allowed us also to reveal an additional lesion tolerance strategy, that we named Damaged Chromatid Loss (DCL), which promotes cell proliferation by replicating only the undamaged chromatid at the expense of losing the damaged chromatid (14) (see Fig. S1C). In the present study, we mainly used the chemical guanine adduct N-2-acetylaminofluorene (G-AAF) positioned in the *Nar*I hotspot sequence: in this configuration the lesion can be bypassed either by the TLS polymerase Pol V in an error-free manner (indicated as TLS0 events), or by the TLS polymerase Pol II, inducing a -2 frameshift (indicated as TLS-2 events) (57).

The *E. coli* strains used are deficient for the mismatch repair system (*mutS*) to prevent correction of the genetic markers of the integrated plasmids, as well as for nucleotide excision repair (*uvrA*), to avoid excision of the lesion and to focus on lesion tolerance events.

### RecBC complex but not RecD participates in lesion bypass

Since RecB is part of the RecBCD complex, we decided to test whether all the subunits of the complex were involved in lesion bypass. We inactivated separately *recB, recC* and *recD* genes and measured tolerance events (*i.e.* HDGR, TLS and DCL) in the presence of a G-AAF lesion. Inactivation of *recB* gene resulted in a decrease of HDGR (from 84% to 61-41% lagging and leading strands, respectively), loss of Pol II TLS events (from 2% to <0.4%) and an increase of DCL events (from 10% to 30%)(see Fig 1), confirming the results previously obtained with UV lesions (14, 20). A *recC* deficient strain behaved like the *recB* strain for lesion tolerance events (Fig 1). On the other hand, no decrease in HDGR or TLS was measured upon inactivation of *recD* (Fig 1), indicating that only the RecBC complex is required for lesion bypass. RecD subunit is involved in the regulation of the nuclease activity of RecB (35, 37). Since we have already shown that the nuclease dead allele of RecB (RecB^D1080A^) has no effect on HDGR or TLS (20), it is not surprising that RecD subunit is not required.

**Figure 1.**
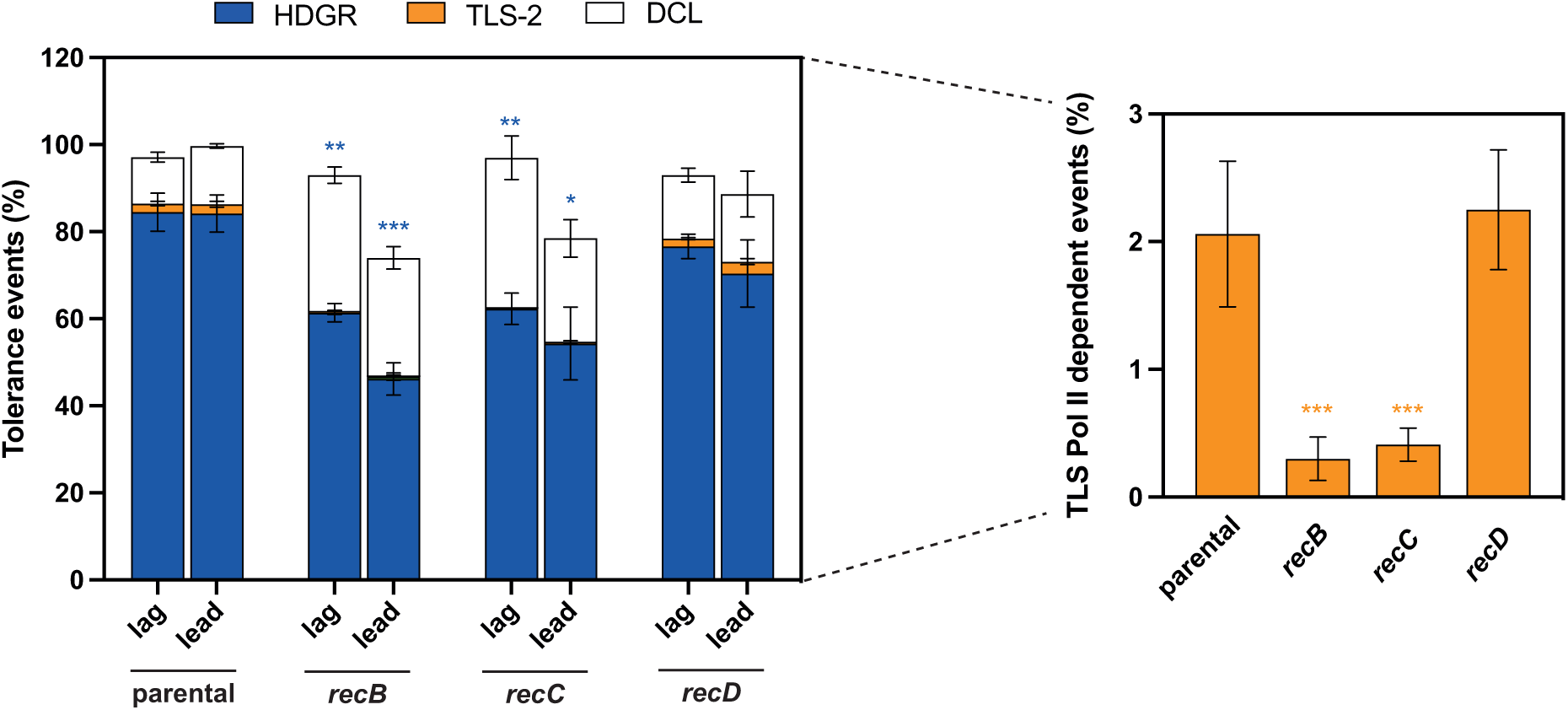
The RecBC complex, without RecD, is involved in lesion bypass. The graph represents the partition of lesion tolerance pathways, that are Homology Directed Gap Repair (HDGR), Translesion Synthesis (TLS) and Damaged Chromatid Loss (DCL), as well as cell survival in the presence of a G-AAF lesion in the mutants of the three subunits of the RecBCD complex (for more details see Material & Methods section). The lesion was inserted either in the lagging (lag) or in the leading (lead) strand. For TLS-2 events, the data for lagging and leading have been pooled together since no difference has been observed. The blue and orange asterisks represent the statistical significance for HDGR and TLS-2 events, respectively, for every mutant strain compared to the parental strain. The data in every graph represent the average and standard deviation of at least three independent experiments. Statistical analyses have been performed with Prism using an unpaired *t*-test. * p < 0,05; ** p < 0,005; *** p < 0,0005. NB: The TLS data for the parental and *recB* strains have been already published in (20).

Further confirmation that RecBC, without RecD, is involved in lesion bypass came using a plasmid construction containing two lesions on opposite strands: this configuration induces a two-fold increase in TLS events due to the generation of overlapping ssDNA gaps that prevent HDGR (16). Deletion of the *recB* gene had a strong effect on TLS levels, while the deletion of *recD* gene or the nuclease dead allele of *recB* behaved in a manner similar to the parental strain (see Fig. S2).

### Exonuclease screen in the *recB* deficient strain

In the absence of RecBC, HDGR mechanism is decreased and TLS is severely impaired. From these data, we hypothesized that RecBC could be involved in the protection of the nascent DNA blocked at the lesion site from nuclease degradation, similarly to what has been described in human cells where BRCA2, together with RAD51, protects the replication fork against the MRE11 nuclease attack (58, 59). Degradation of the 3’ end of the nascent DNA is detrimental for TLS polymerases since their cognate substrate (*i.e.* the 3’ end blocked at the lesion site) is missing, and uncontrolled nascent DNA resection may also impact the efficiency of HDGR mechanism. To verify our hypothesis, we performed a genetic screen to identify the nuclease(s) potentially responsible for this degradation in the absence of *recB*. We looked for candidate genes that encode exonucleases with a 3’->5’ polarity that preferentially degrade single-stranded DNA and that have a role in DNA metabolism and genome maintenance. Based on the literature (see reviews (60, 61)), we tested ExoI (also known as XonA/SbcB), ExoVII, ExoX and SbcCD.

We inactivated each exonuclease individually in the *recB* deficient strain and measured Pol II dependent TLS events in the presence of a G-AAF lesion in order to see whether we could restore the level of the parental strain. From this first screen, ExoI and SbcCD emerged as good candidates, since deletion of *exoI* or *sbcCD* in the *recB* strain recovered fully or partially the level of TLS-2 (Fig 2A).

**Figure 2.**
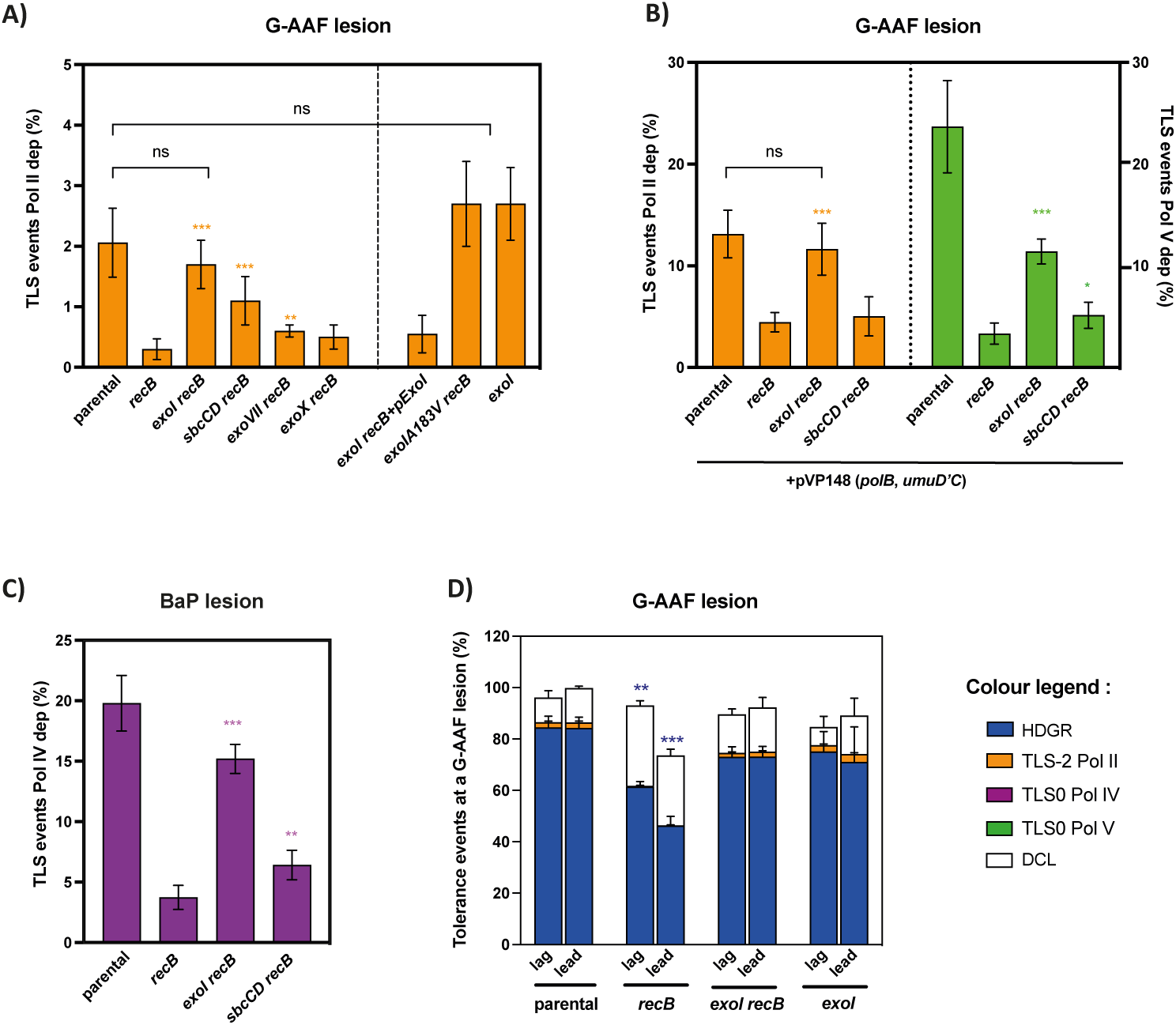
Exonuclease screen. **A)** The graph represents the percentage of Pol II TLS events in the presence of a G-AAF lesion for several exonuclease mutant strains combined with *recB* deletion. The orange asterisks represent the statistical significance for every exonuclease mutant strain compared to *recB* deficient strain. pExoI is a plasmid expressing *exoI* gene under its own promoter. ExoIA183V is the nuclease dead allele of ExoI (also known as *sbcB15*). **B)** The graph represents the percentage of Pol II and Pol V TLS events in the presence of a G-AAF lesion. All the strains contain the pVP148 plasmid that expresses *polB* and *umuD’C* genes under their native promoters. The orange and green asterisks represent the statistical significance for the mutant strains compared to *recB* deficient strain. **C)** The graph represents the percentage of Pol IV TLS events in the presence of a BaP lesion. The violet asterisks represent the statistical significance for the mutant strains compared to *recB* deficient strain. The data in every graph represent the average and standard deviation of more than four independent experiments obtained in the lagging or in the leading orientation. Statistical analyses have been performed with Prism using an unpaired *t*-test: * p < 0,05; ** p < 0,005; *** p < 0,0005. **D)** The graph represents the partition of lesion tolerance pathways (*i.e.* HDGR, TLS and DCL) and cell survival in the presence of a G-AAF lesion for *exoI* deletion strains. The lesion was inserted either in the lagging (lag) or in the leading (lead) strand. The data represent the average and standard deviation of at least three independent experiments. No statistically difference for the *exoI* strains (unpaired *t-*test analysis) was observed compared to the parental strain. NB: The data for the parental and *recB* strains in A), C) and D) panels have been already published in (20).

### ExoI prevents efficient lesion bypass in the absence of RecBC

We have previously shown that the deletion of *recB* strongly affects lesion bypass by all three TLS polymerases (20). As mentioned above, when positioned in the *Nar*I hotspot sequence, the G-AAF lesion can be bypassed by Pol II, but also by Pol V. However, under physiological conditions, Pol V bypass is very low (<0.5%) (13). Hence, to assess the role of ExoI and SbcCD on Pol V TLS events, we used a plasmid expressing both the active form of Pol V (UmuD’C in which UmuD is already cleaved (54, 62)) and Pol II under their own promoters (15). The expression level of both TLS polymerases is 3-5 times higher than in the parental strain, which results in an increase of Pol II TLS events (from 2% to 12%) and of Pol V TLS events (from 0.5% to more than 20%) (Fig 2B). These conditions allow a better dynamic range to monitor the effect of the nucleases on both Pol V and Pol II-mediated TLS. As expected, deletion of the *recB* gene strongly decreased the level of both Pol II and Pol V dependent TLS events (Fig 2B). While deletion of the nuclease *sbcCD* in the *recB* strain did not restore TLS levels, the *exoI recB* double mutant fully restored Pol II-dependent TLS events and partially restored Pol V-dependent TLS events (Fig 2B).

Finally, we tested the effect of the two nucleases on the TLS polymerase Pol IV in the presence of the benzo(a)pyrene (BaP) lesion (63). Only the deletion of *exoI* in the *recB* deficient strain was able to almost fully restore the TLS level of the parental strain (Fig 2C), thus pointing ExoI as the major nuclease acting in the absence of RecBC.

Since loss of RecBC also affects HDGR mechanism (Fig 1), we measured the HDGR level in an *exoI recB* strain in the presence of a G-AAF lesion and observed a full recovery of the HDGR level, similar to the parental strain (Fig 2D). Hence, from these data it appears that the defect in lesion bypass observed in the absence of RecBC can be largely alleviated by inactivation of ExoI.

### RecBC protects from ExoI exonuclease

ExoI is a Mg^2+^-dependent exonuclease that rapidly processes ssDNA in the 3’->5’ direction (64). It is a highly processive nuclease whose activity is stimulated by the single stranded DNA binding protein (SSB) (65). ExoI plays several roles in DNA metabolism and DNA repair, in particular in mismatch repair, recombination and completion of DNA replication (23, 66–68). Hence, we pursued the characterization of the role of ExoI in lesion bypass. As expected, complementation experiments in the *exoI recB* strain, using a plasmid carrying a functional *exoI* gene under its native promoter, resulted in low levels of Pol II TLS events in the presence of a G-AAF lesion similar to the one observed in a *recB* deficient strain (Fig 2A: *exoI recB*+p*ExoI*).

To confirm our hypothesis that it is the exonuclease activity of ExoI that is preventing efficient lesion bypass in the absence of RecBC, we used the well-described mutant allele *sbcB15* (*exoIA183V*), which is inactivated in its nuclease active site (68). We introduced the point mutation in the *exoI* gene by CRISPR-Cas9 technology in the *recB* deficient strain and measured the level of Pol II TLS in the presence of a G-AAF lesion. We observed full recovery of Pol II TLS, similar to the parental strain (Fig 2A: *exoIA183V recB*)).

Taken together these data indicate that in the absence of RecBC, ExoI is the major exonuclease that negatively affects lesion bypass. Given that its exonuclease catalytic activity is required, we surmise that it acts by degrading the nascent DNA that presents a 3’ extremity blocked at the lesion site. We wondered whether in the presence of RecBC, ExoI could still access to the 3’ end of the nascent DNA, thus affecting lesion bypass. We inactivated *exoI* gene in the parental strain and measured Pol II TLS events and HDGR in the presence of a G-AAF lesion. No change in the level of Pol II TLS events, nor in HDGR events were observed compared to the parental strain (Fig 2A and 2D) suggesting that in the presence of RecBC, ExoI has no effect on lesion bypass.

### ExoI and the TLS polymerases are in competition for the same substrate

Based on these genetic data, we propose that the RecBC complex protects the nascent DNA blocked at the lesion site from the attack of the ExoI nuclease, in order to allow efficient lesion bypass, either by TLS polymerases or by HDGR mechanism. In the absence of RecBC, ExoI can access the 3’ end of the nascent DNA, resecting it and affecting lesion bypass. According to this model, in the absence of RecBC, the action of ExoI would be in competition with lesion tolerance pathways, in particular with the TLS polymerases since their cognate substrate (the 3’ end) is the same. Therefore, overexpression of the TLS polymerases could partially compensate for the absence of RecBC. Indeed, in a *recB* strain, we can see that TLS events are much higher when Pol II and Pol V are overexpressed compared to the same strain with basal level of Pol II and Pol V (compare Fig 2A to 2B). Further confirmation came comparing Pol II TLS levels in a *lexA* deficient strain in the presence or absence of RecB: in such strain, constitutive SOS induction leads to a strong increase in the expression level of TLS polymerases. In a *recB lexA* strain, Pol II dependent TLS is much less affected compared to a *recB* strain, indicating that overexpression of Pol II can partially compensate for RecBC absence (Fig. S3).

These data suggest that ExoI, whose substrate is the nascent 3’ DNA end at the lesion site, is in competition with the TLS Polymerases that share the same substrate.

### RecBC is required in the presence of strong blocking lesions

We have shown the requirement for RecBC in the presence of strong blocking lesions such as G-AAF and UV lesions (14, 20), when a ssDNA gap is formed downstream of the lesion. When the replisome encounters a less blocking lesion, we can expect that the bypass could occur without the need for replication to restart downstream of the lesion: in this situation, no ssDNA gap will form as the lesion will be rapidly bypassed by the replicative polymerase or a TLS polymerase. Therefore, we asked whether in this situation RecBC was still required to protect the nascent DNA. For this purpose, we used three less blocking lesions, the amino fluorene (AF) (69), the hydroxypropyl (1hpG) (70) and the N2-G-furfuryl (18). As shown in Fig. 3, TLS events at these lesions are higher than for the strong blocking lesions such as the G-AAF or the UV photoproducts (13). In the absence of RecBC, a less than two-fold decrease in lesion bypass was observed for the three lesions (Fig. 3), while we have measured a 5 to 7-fold decrease in the absence of RecBC when stronger blocking lesions were used (see (20) and Fig. 2). These data indicated that RecBC requirement is more critical in the presence of a persistent ssDNA gap: when the 3’ end of the nascent DNA is exposed longer, its protection becomes more critical to allow TLS and HDGR mechanisms.

**Figure 3:**
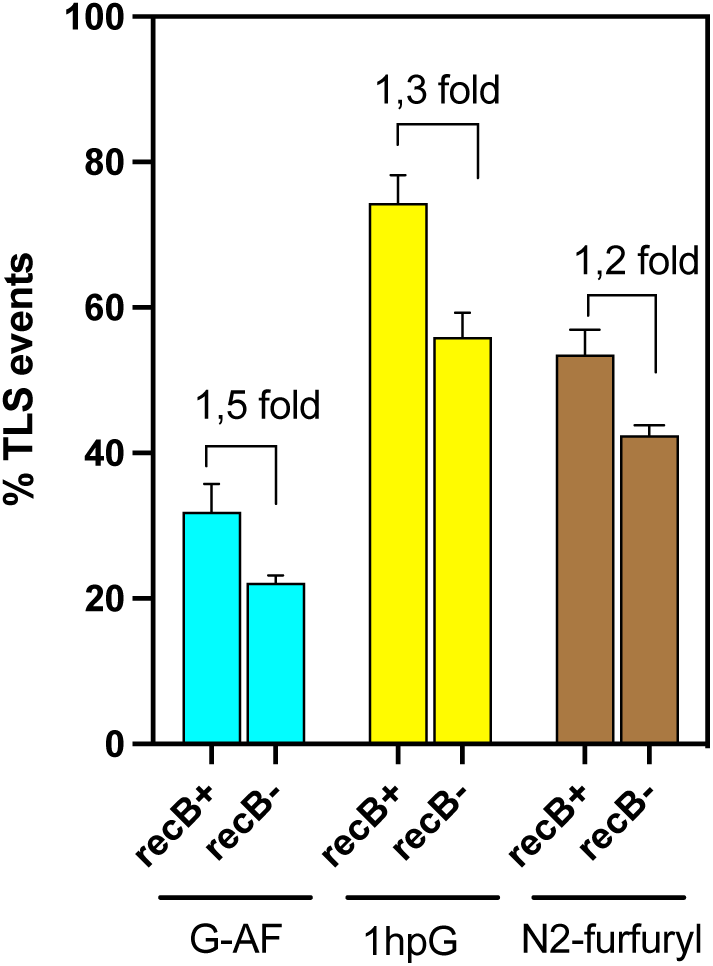
RecBC is required for efficient lesion bypass only in the presence of strong-replication blocking lesions. The graph represents the tolerance events TLS like in the presence of less blocking lesions due to DNA pol III or to a TLS polymerase bypass. The data represent the average and standard deviation of more than four independent experiments obtained in the lagging or in the leading orientation.

### RecBC complex binds to ssDNA gap in the presence of SSB

Our genetic data indicate that the nascent 3’ DNA end is somehow protected by the RecBC complex and that this protection is more critical for persistent ssDNA gaps. Since the protective role of RecBC is independent of its enzymatic activities (20), we surmise that a structural action could occur by the binding of RecBC to the ssDNA gap. In order to test this hypothesis, we performed *in vitro* assays to assess whether RecBC was indeed binding to a ssDNA gap.

For this purpose, we expressed and purified the RecBC complex according to (38) (see Fig. S4A). As mentioned before, the RecBC complex was shown to lack exonuclease activity while still preserving a weak helicase activity (48, 71, 72). Our purified RecBC complex was indeed not able to degrade a linear blunt DNA substrate (Fig. S4B) but we observed a weak helicase activity with the same DNA substrate (Fig. S4C). We also confirmed that our purified RecBC complex was able to bind to a double stranded substrate (Fig. S4D) as shown by (73).

In parallel, we constructed two gapped DNA substrates: i) a 5.3 kb gap consisting of the phage phiX174 (ϕX174) to which is hybridized a 50 nt radiolabelled oligonucleotide; ii) a 2.5 kb gap consisting of the phage ϕX174 to which is hybridized a complementary 2.8 kb ssDNA (Fig. 4A). We then performed electrophoretic mobility shift assays using the 5.3 kb gapped substrate and our purified RecBC complex. When using the RecBC complex alone, no shift was observed (Fig. 4B). We reasoned that once a ssDNA gap is produced *in vivo*, it is rapidly covered by the single-stranded binding protein SSB. Therefore, we tested whether the presence of SSB could allow the binding of RecBC to the ssDNA gap. First, we tested different concentrations of SSB that allowed covering the entire ssDNA gap (Fig. S5A). We chose three concentrations of SSB and added increasing concentration of RecBC: interestingly RecBC was able to bind to a SSB-covered ssDNA gap in a concentration-dependent manner (Fig. 4C). The strongest shift was obtained with a high concentration of RecBC (85nM) that corresponded to a RecBC/DNA ratio of roughly 300-400:1. We decided to keep this protein/DNA ratio for RecBC and test different concentrations of SSB: a shift was again observed for the different concentrations of SSB, until we reached a high concentration of SSB where RecBC became unable to bind to the substrate (Fig. S5B). We obtained similar results with the 2.5 kb gap construction (Fig. S5C). These results show that RecBC binds to a SSB-covered ssDNA gap, unless SSB concentration is too high suggesting that a certain mobility in SSB binding is required to allow RecBC binding.

**Figure 4:**
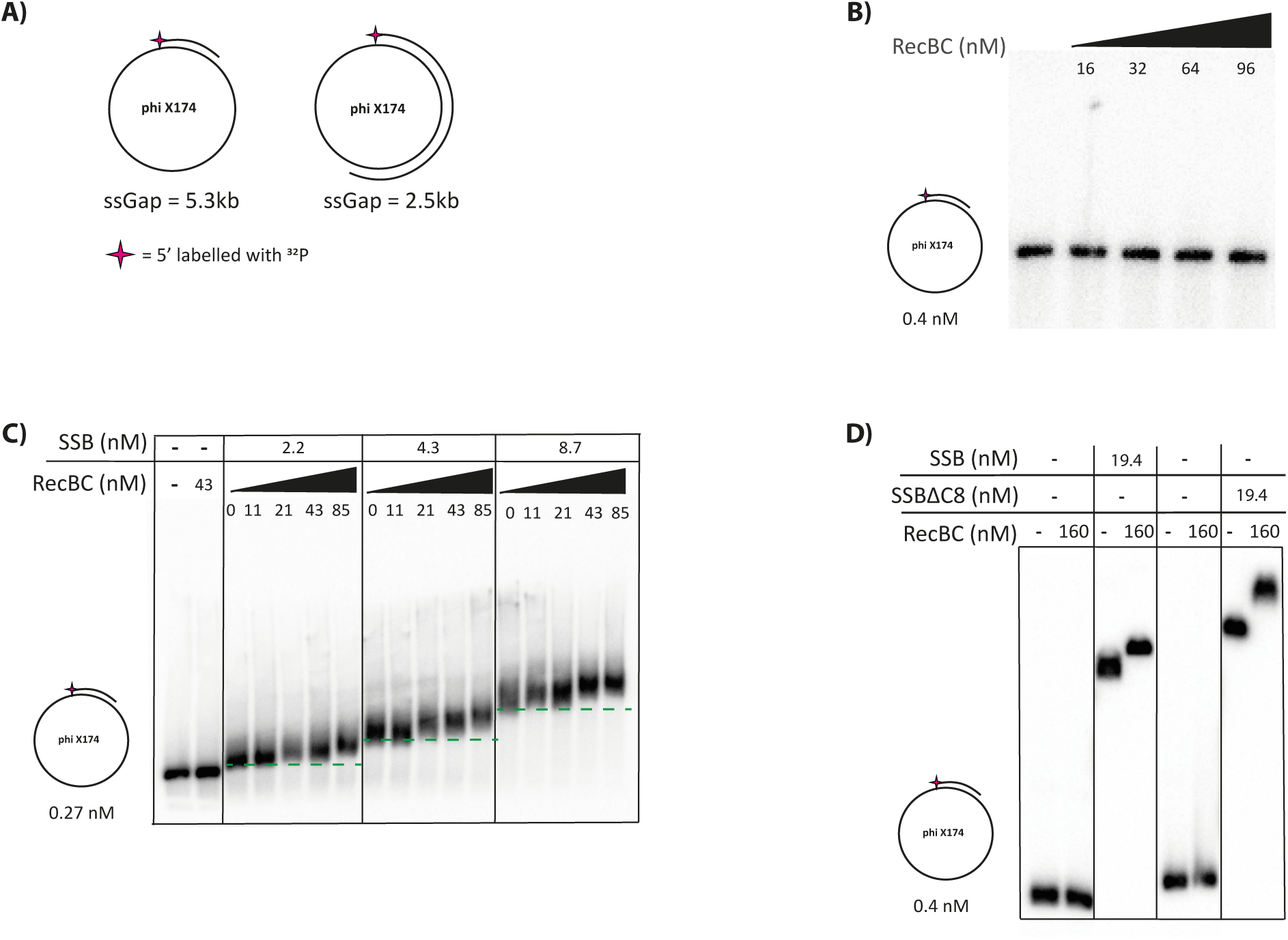
RecBC complex binds to a ssDNA gap in the presence of SSB. **A)** DNA substrates used for the electrophoretic mobility shift assays. The circular ssDNA of the phage ϕX174 was used to obtained a 5.3 kb ssDNA gap or a 2.5 kb ssDNA gap. The 5’of the ds-ss junction of the substrates have been radiolabeled with ^32^P. **B)** Mobility shift assays on agarose gel using the 5.3 kb gapped substrate in the presence of increasing concentrations of RecBC complex. **C)** Mobility shift assay on agarose gel using the 5.3 kb DNA gap in the presence of three non-saturating concentrations of SSB and increasing concentrations of RecBC complex (for more details see Materials and Methods). **D)** Mobility shift assay on agarose gel using the 5.3 kb DNA gap in the presence of the truncated form of SSB (SSBΔ8).

### The acidic tip of SSB is not required for RecBC binding to DNA gaps

To characterize the cooperative binding of RecBC with SSB, we asked whether there was a direct interaction between RecBC and SSB. Indeed, bacterial SSBs are known to be a hub for protein recruitment, in particular their C-terminal tail contains several amino acids (also called the “acidic tip”) responsible for the binding of several proteins involved in recombination, repair and replication (74–79). We performed mobility shift assay, using a well-described truncated form of SSB (SSBΔ8) that lacks the last 8 amino acids, thus affecting most of protein-protein interactions (80), but not the DNA binding capacity (74). In the presence of the SSBΔ8, the RecBC complex was still able to bind to the substrate (Fig. 4D), suggesting no direct interaction between the two proteins via this acidic tip. However, we cannot rule out the possibility that RecBC does interact with SSB through another region, like for instance its intrinsically disordered linker (IDL) domain since it has been shown recently that this part of the protein might also be involved in protein-protein interactions (78, 79, 81).

### ExoI needs a helicase to access the nascent DNA

ExoI is known to digest single stranded DNA in the 3’->5’ direction with high processivity and to dissociate upon encountering with dsDNA, unless a helicase provides the right substrate (61, 64). In agreement with this, when we tested the activity of ExoI on our gapped DNA substrate, we did not observe any degradation except when we added a flap of 6 nt or 12 nt (Fig. S6A). The flap was degraded up to 2 nt before the ds-ss DNA junction, as expected since ExoI cannot degrade further. Therefore, we hypothesize that ExoI acts on the nascent DNA in combination with a helicase. In an attempt to identify this putative helicase, we have performed a helicase screen (only non-essential helicases could be tested) in the *recB* deficient strain, similar to the one done for the nucleases. None of the tested candidates (*uvrD, dinG, yoaA, helD* genes) restored the level of TLS as the parental strain (Fig. S6B). Without the identification of this helicase, we were not able to assess *in vitro* that the presence of RecBC prevents degradation of the 3’ end of the gapped substrate. However, we observed that RecBC prevents ExoI action when RecBC is bound to an oligo (Fig. S6C), suggesting that this could also occur when RecBC binds to a ssDNA gap.

### RecBC binds internally to the SSB-covered ssDNA gap

In order to understand how RecBC binds and protects the ssDNA gaps, we analyzed the DNA-protein complexes formed by the 2.5 kb gapped DNA substrate and purified RecBC in presence or not of SSB by using transmission electron microscopy (TEM). We first incubated the DNA gapped substrate in the absence of protein, and observed that the ssDNA region folded into secondary structures (Fig. 5A and Fig. S7A). In agreement with the EMSA experiment, RecBC was not able to bind to a naked gapped substrate in the absence of SSB (Fig. S7B). We then incubated the gapped substrate with SSB protein, using similar conditions to the *in vitro* shift assays: this resulted in SSB partially covering the ssDNA gap and creating a nicely folded SSB filament (Fig. 5B-C). In the presence of SSB the addition of RecBC, at the concentration used for the gel shift assay, induced a more compacted and tangled structure of the SSB-covered ssDNA substrate (Fig 5D-E-F). We quantified this RecBC compaction effect by i) measuring the SSB-ssDNA contour length (Fig 5G) and ii) the distance between the two ds-ss junctions (Fig. S7H). Both measuring methods showed that the binding of RecBC significantly reduced the SSB-ssDNA length. The same compaction was observed and quantified in presence of a lower SSB concentration (Fig. S7C-D versus Fig. S7E-F-G) confirming our EMSA data (Fig. 4C). The TEM analysis suggests that RecBC oligomerizes and could bind internally to the SSB-covered ssDNA gap but also to the ds-ss junction (Fig 5D-E-F, see white arrows). Previous studies have already showed SSB ability to diffuse along the DNA thus releasing segments of ssDNA, and providing access to other proteins, such as RecA (reviewed in (75, 78)). The binding of RecBC is clearly specific of a SSB-covered gapped substrate since no RecBC binding was observed in the presence of the SSB-covered circular ssDNA ϕX174 (Fig. S7K). This data also supports the idea that RecBC does not directly interact with SSB, as already suggested by the *in vitro* shift assays, but rather needs a ss-dsDNA junction to load onto the substrate.

**Figure 5:**
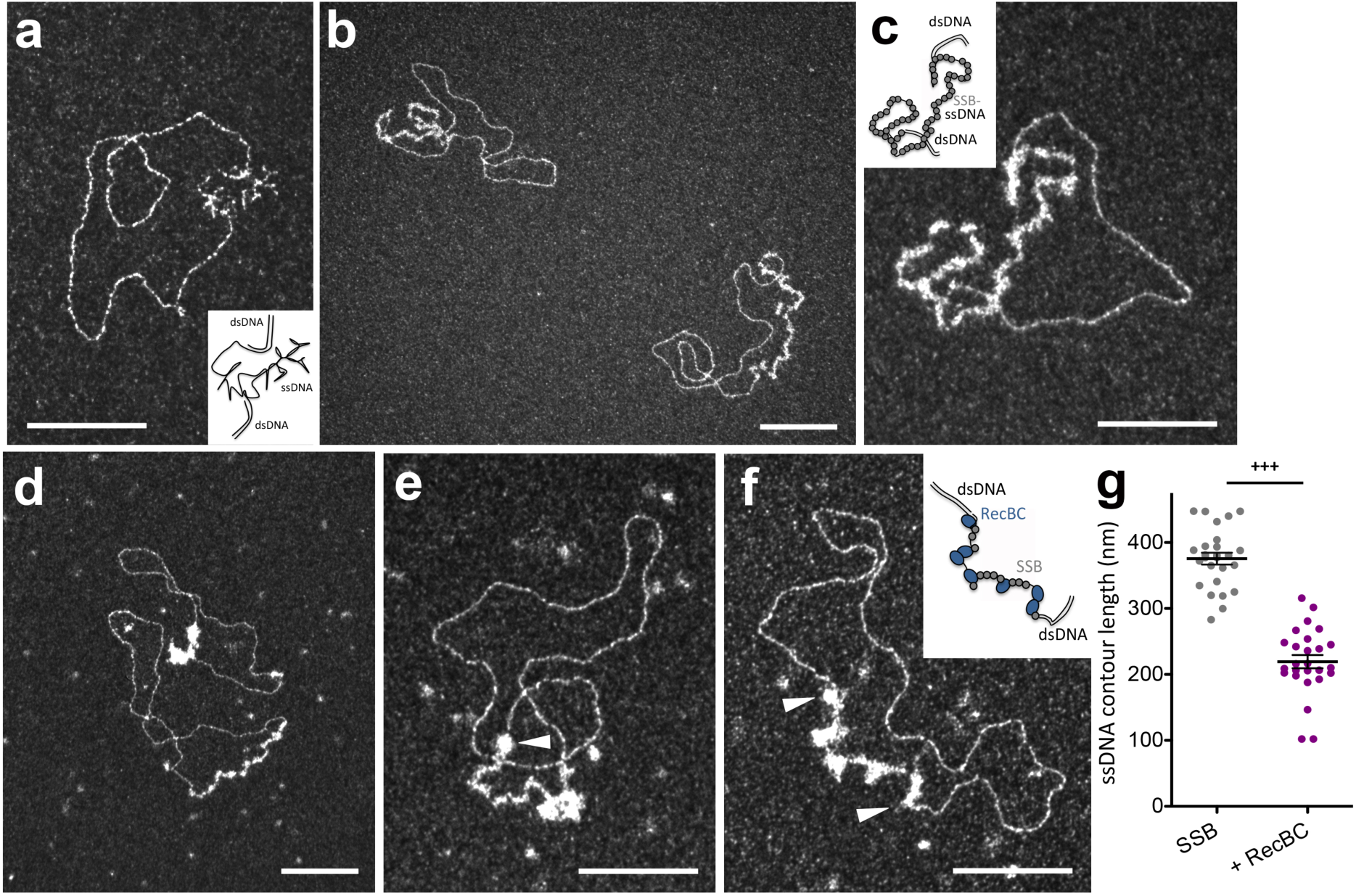
Transmission Electron Microscopy analysis of RecBC-SSB-ssDNA gap complexes in darkfield imaging mode. **a)** Control DNA substrate containing a 2.5 kb ssDNA gap. ssDNA is folded into secondary structures under these spreading conditions. **b-c**) 286 nM SSB were incubated with the 2.5 kb ssDNA gapped substrate (10 μM nucleotides - 3.2 nM in molecules). The ssDNA part of the substrate is partially covered by SSB protein. **d-f)** 200 nM of RecBC were added to the reaction. RecBC binding to SSB-ssDNA induces its compaction. White arrows indicate RecBC oligomers localization to the ss-dsDNA junctions. The scale bars correspond to 100 nm. **g)** Distribution of the SSB-ssDNA contour length (in nm) in absence (grey points) or in presence of 200 nM RecBC (magenta points). The statistical unpaired t-test was applied, *** corresponded to a P value < 0.001.

The binding of RecBC to the ssDNA gap generates compacted and entangled structures that hinder the access of ExoI and/or the putative helicase. Consequently, RecBC binding protects the nascent DNA by blocking helicase and nuclease activity.

## DISCUSSION

This study unveils a cellular role for the RecBC complex in protecting the nascent DNA to allow efficient lesion bypass in the presence of a single-stranded DNA gap. Indeed, in the absence of RecBC lesion bypass is severely compromised, because degradation of the nascent DNA by the nuclease ExoI prevents the action of the TLS polymerases and also affects the efficiency of the HDGR mechanism. While we have recently shown that the ssDNA gap needs to be enlarged on its 5’ end by the action of the nuclease RecJ and the helicase RecQ before the mediator proteins recF-O-R load RecA (17), the present study shows that the 3’ end of the gap needs to be protected from nucleolytic degradation operated by the exonuclease ExoI to enable efficient lesion bypass and complete replication.

### RecBC protection is required in the presence of persistent ssDNA gaps

Based on our *in vivo* and *in vitro* results, we propose a model (Fig. 6) in which RecBC is recruited to a persistent SSB-covered ssDNA gap. This situation is typical of a strong replication-blocking lesion when replication needs to restart downstream in order to continue replication, creating a ssDNA gap that will be later filled in by one of the two lesion tolerance pathways. The ssDNA gap is immediately covered by the SSB protein that protects the ssDNA and orchestrates the recruitment of several proteins to preserve genome integrity. We propose that RecBC complex(es) binds to the “naked” (*i.e.* SSB-free) ssDNA sections left available by the dynamic movement of the SSB tetramers along the ssDNA gap. The binding of RecBC results in a more compacted and entangled structure of the SSB filament, and this structure will indirectly prevent the action of the nuclease ExoI (together with an unidentified helicase), protecting the nascent DNA and allowing an efficient lesion bypass. On the other hand, when replication is stalled by a less blocking lesion that can be easily bypassed by either the replicative polymerase or a TLS polymerase, the ssDNA created by the progression of the replicative helicase is only transient/limited and RecBC protection is not as critical.

**Figure 6:**
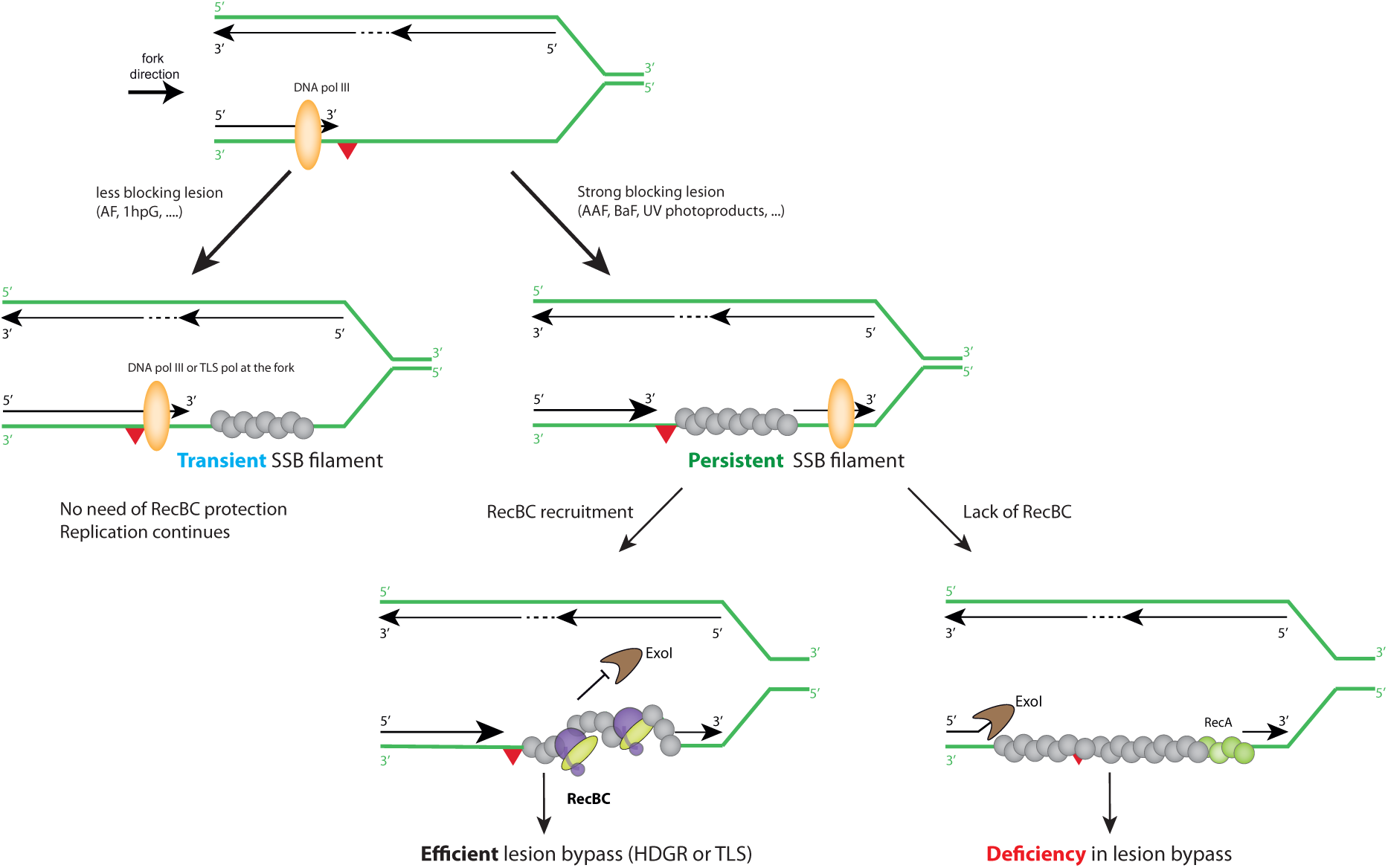
Model of RecBC complex protecting the nascent DNA in the presence of a ssDNA gap. The replicative DNA polymerase is temporarily blocked at a DNA lesion : i) when the lesion can be easily bypassed by the replicative DNA polymerase itself or by a TLS polymerase at the fork, the SSB-covered ssDNA formed by the DNA unwinding is transient and the RecBC complex is not needed for protection; ii) in the presence of a strong blocking lesion, replication restarts downstream creating a persistent SSB-covered ssDNA gap: RecBC is recruited and binds to the SSB-covered ssDNA gap by restructuring and condensing the ssDNA to protect the nascent DNA against nuclease attack. This will allow efficient lesion bypass by one of the two lesion tolerance pathways. In the absence of RecBC, ExoI can access the 3’ end of the nascent DNA (potentially with the help of a helicase) degrading the nascent DNA. This will impact lesion bypass, in particular the action of the TLS polymerases, but also the HDGR mechanism.

SSB is a constitutive component of the replisome, associating with the ssDNA on the lagging strand during Okazaki fragment formation or ssDNA created during the DNA helicase unwinding. However, the SSB binding lifetime to these “physiological” ssDNA gaps and the SSB binding lifetime to the ssDNA gaps formed upon lesion encountering are probably different, as recently suggested by Loparo’s group (82). We also showed with the same group that the mobility change in SSB molecules upon replication perturbation could act as a molecular switch to enrich proteins at the lesion site, as it is the case for Pol IV TLS polymerase (18, 82). Hence, such changes in SSB dynamics support our model in which RecBC is specifically recruited to protect the nascent DNA of the ssDNA gap formed in the presence of strong blocking lesions and not in the presence of less blocking lesions that generate more transient gaps. Our *in vitro* data (both the gel shift assays and the TEM images) further support this idea, as RecBC binding is observed only in the presence of an SSB-covered gapped substrate, reinforcing the notion that SSB dynamics play a key role in RecBC recruitment.

### Degradation of nascent DNA strongly affects lesion tolerance pathways

In the absence of RecBC, the SSB filament folds properly onto the ssDNA gap allowing for ExoI to access the nascent DNA with the help of a yet-to-be identified helicase. ExoI will then degrade the nascent DNA, thus strongly inhibiting TLS polymerases access/activity. Therefore, in the absence of RecBC, ExoI and the TLS polymerases are in competition for the same substrate (the 3’ end). In line with this, we showed that overexpression of the TLS polymerases can partially outcompete the action of ExoI in a *recB* deficient strain. Undesired degradation of the nascent DNA also affects HDGR mechanism possibly because more RecA molecules will be necessary to form the recombinogenic filament and subsequently more time for the ssDNA gap to be filled will be needed, leading to an increase in damage chromatid loss events (DCL, Fig 1). However, high levels of HDGR still persist in the absence of RecBC, suggesting that the degradation of the nascent DNA by ExoI is limited, or that the action of ExoI is also kinetically in competition with the RecA filament formation. Indeed, it is likely that once the recombinogenic filament is formed, it will engage in homology search and strand invasion and ExoI will not be able to access the nascent DNA. This is corroborated by the phenotype of the double mutant *recF recB* in our previous study (14) where we observed a strong loss of HDGR and cell survival (similar to a *recA* deficient strain). Delaying the formation of the RecA filament (in the absence of its mediator RecF) in addition to the loss of RecBC will favor ExoI activity that will degrade the nascent DNA and therefore strongly impact HDGR mechanism and cell survival.

In the last decade, growing evidence has indicated that the nascent DNA must be protected against uncontrolled and excessive nucleolytic degradation. Failure to do so can severely compromise DNA replication and genome stability (83, 84). Surprisingly, proteins usually involved in homologous recombination pathway, such as RAD51 and BRCA1/2 in humans, came to the fore as guardians of the nascent DNA against the attack of specific nucleases (59, 85, 86). This function is distinct from their canonical function in homologous recombination. Similarly, in this study we have shown that RecBC, that is part of the RecBCD complex involved in homologous recombination repair in *E. coli* and is one of the mediators of RecA, plays a new and unexpected role in lesion bypass by protecting the replication fork. This role is independent of its canonical catalytic activities. This new finding indicates once more how the mechanisms of DNA replication are evolutionary conserved in all living organisms. It will be interesting to investigate in human cells whether lesion bypass may also be compromised in cells deficient for the so-called fork protection when a ssDNA gap is formed.

## DATA AVAILABILITY

All relevant data are within the manuscript or in the Supplementary files and are available on request.

## FUNDING

This work was supported by Fondation pour la Recherche Médicale [Equipe FRM-EQU201903007797] https://www.frm.org (VP), by ANR (17-CE12-0015) and ERM-Université Paris-Saclay (ELC and PD). Funding for open access charge: CNRS. The funders had no role in study design, data collection and analysis, decision to publish, or preparation of the manuscript.

## ACKNOWLEDGEMENTS

We thank Jean-Hugues Guervilly for critical reading of the manuscript. We thank Dave Wigley for kindly providing us the plasmids for RecBC expression/purification and Michael Cox for kindly providing us the SSBΔ8 protein.

## Notes

### Competing Interest Statement

The authors have declared no competing interest.

### Summary of Updates

Rewriting some paragraphs, more Electron Microscopy data

## REFERENCES

1. Pagès V, Fuchs RP (2003) Uncoupling of leading- and lagging-strand DNA replication during lesion bypass in vivo. *Science (New York*, NY) 300(5623):1300–1303.

2. Yeeles JTP, Marians KJ (2013) Dynamics of leading-strand lesion skipping by the replisome. Molecular Cell 52(6):855–865.

3. Beattie TR et al. (2017) Frequent exchange of the DNA polymerase during bacterial chromosome replication. Elife 6:e21763.

4. Yang W, Seidman MM, Rupp WD, Gao Y (2019) Replisome structure suggests mechanism for continuous fork progression and post-replication repair. DNA Repair (Amst) 81:102658.

5. Heller RC, Marians KJ (2006) Replication fork reactivation downstream of a blocked nascent leading strand. Nature 439(7076):557–562.

6. Yeeles JTP, Marians KJ (2011) The Escherichia coli replisome is inherently DNA damage tolerant. Science (New York, NY) 334(6053):235–238.

7. Henrikus SS et al. (2018) DNA polymerase IV primarily operates outside of DNA replication forks in Escherichia coli. PLoS genetics 14(1):e1007161.

8. Quinet A, Tirman S, Cybulla E, Meroni A, Vindigni A (2021) To skip or not to skip: choosing repriming to tolerate DNA damage. Mol Cell

9. Pham P, Shao Y, Cox MM, Goodman MF (2022) Genomic landscape of single-stranded DNA gapped intermediates in Escherichia coli. Nucleic Acids Res 50(2):937–951.

10. Branzei D, Psakhye I (2016) DNA damage tolerance. Current opinion in cell biology 40:137–144.

11. Henrikus SS, van Oijen AM, Robinson A (2018) Specialised DNA polymerases in Escherichia coli: roles within multiple pathways. Current genetics 64(6):1189–1196.

12. Ashour ME, Mosammaparast N (2021) Mechanisms of damage tolerance and repair during DNA replication. Nucleic Acids Research 49(6):3033–3047.

13. Pagès V, Mazón G, Naiman K, Philippin G, Fuchs RP (2012) Monitoring bypass of single replication-blocking lesions by damage avoidance in the Escherichia coli chromosome. Nucleic acids research 40(18):9036–9043.

14. Laureti L, Demol J, Fuchs RP, Pagès V (2015) Bacterial proliferation: keep dividing and don’t mind the gap. PLoS genetics 11(12):e1005757.

15. Naiman K, Philippin G, Fuchs RP, Pagès V (2014) Chronology in lesion tolerance gives priority to genetic variability. Proceedings of the National Academy of Sciences of the United States of America 111(15):5526–5531.

16. Chrabaszcz É, Laureti L, Pagès V (2018) DNA lesions proximity modulates damage tolerance pathways in Escherichia coli. Nucleic acids research 46(8):4004–4012.

17. Laureti L, Lee L, Philippin G, Kahi M, Pagès V (2022) Single strand gap repair: The presynaptic phase plays a pivotal role in modulating lesion tolerance pathways. PLoS Genet 18(6):e1010238.

18. Chang S et al. (2022) Compartmentalization of the replication fork by single-stranded DNA-binding protein regulates translesion synthesis. Nat Struct Mol Biol 29(9):932– 941.

19. Pagès V (2016) Single-strand gap repair involves both RecF and RecBCD pathways. Current genetics

20. Laureti L, Lee L, Philippin G, Pagès V (2017) A non-catalytic role of RecBCD in homology directed gap repair and translesion synthesis. Nucleic Acids Research 45(10):5877–5886.

21. Arnold DA, Kowalczykowski SC (2001) RecBCD Helicase/Nuclease. Encyclopedia of life sciences 1–6.

22. Dillingham MS, Kowalczykowski SC (2008) RecBCD enzyme and the repair of double-stranded DNA breaks. Microbiology and molecular biology reviews : MMBR 72(4):642–671.

23. Wendel BM, Courcelle CT, Courcelle J (2014) Completion of DNA replication in Escherichia coli. Proceedings of the National Academy of Sciences 111(46):16454– 16459.

24. Kowalczykowski SC, Dixon DA, Eggleston AK, Lauder SD, Rehrauer WM (1994) Biochemistry of homologous recombination in Escherichia coli. Microbiological reviews 58(3):401–465.

25. Kuzminov A (1999) Recombinational repair of DNA damage in Escherichia coli and bacteriophage lambda. Microbiology and molecular biology reviews : MMBR 63(4):751–813.

26. Spies M, Kowalczykowski SC (2005) Homologous recombination by RecBCD and RecF pathways. The bacterial chromosome 389–403.

27. Amundsen SK, Smith GR (2023) RecBCD enzyme: mechanistic insights from mutants of a complex helicase-nuclease. Microbiol Mol Biol Rev e0004123.

28. Hickson ID, Robson CN, Atkinson KE, Hutton L, Emmerson PT (1985) Reconstitution of RecBC DNase activity from purified Escherichia coli RecB and RecC proteins. Journal of Biological Chemistry 260(2):1224–1229.

29. Masterson C et al. (1992) Reconstitution of the activities of the RecBCD holoenzyme of Escherichia coli from the purified subunits. Journal of Biological Chemistry 267(19):13564–13572.

30. Handa N, Ohashi S, Kusano K, Kobayashi I (1997) Chi-star, a chi-related 11-mer sequence partially active in an E. coli recC1004 strain. Genes Cells 2(8):525–536.

31. Arnold DA, Handa N, Kobayashi I, Kowalczykowski SC (2000) A novel, 11 nucleotide variant of chi, chi*: one of a class of sequences defining the Escherichia coli recombination hotspot chi. J Mol Biol 300(3):469–479.

32. Handa N et al. (2012) Molecular determinants responsible for recognition of the single-stranded DNA regulatory sequence, χ, by RecBCD enzyme. Proc Natl Acad Sci U S A 109(23):8901–8906.

33. Amundsen SK, Neiman AM, Thibodeaux SM, Smith GR (1990) Genetic dissection of the biochemical activities of RecBCD enzyme. Genetics 126(1):25–40.

34. Chamberlin M, Julin DA (1996) Interactions of the RecBCD Enzyme from Escherichia coliand Its Subunits with DNA, Elucidated from the Kinetics of ATP and DNA Hydrolysis with Oligothymidine Substrates †. Biochemistry 35(50):15949–15961.

35. Yu M, Souaya J, Julin DA (1998) Identification of the nuclease active site in the multifunctional RecBCD enzyme by creation of a chimeric enzyme. Journal of Molecular Biology 283(4):797–808.

36. Churchill JJ, Anderson DG, Kowalczykowski SC (1999) The RecBC enzyme loads RecA protein onto ssDNA asymmetrically and independently of χ, resulting in constitutive recombination activation. Genes & development 13(7):901–911.

37. Amundsen SK, Taylor AF, Smith GR (2000) The RecD subunit of the Escherichia coli RecBCD enzyme inhibits RecA loading, homologous recombination, and DNA repair. Proceedings of the National Academy of Sciences of the United States of America 97(13):7399–7404.

38. Wilkinson M, Chaban Y, Wigley DB (2016) Mechanism for nuclease regulation in RecBCD. Elife 5:e18227.

39. Taylor AF, Smith GR (1985) Substrate specificity of the DNA unwinding activity of the RecBC enzyme of Escherichia coli. Journal of Molecular Biology 185(2):431–443.

40. Taylor AF, Smith GR (1995) Monomeric RecBCD Enzyme Binds and Unwinds DNA (∗). Journal of Biological Chemistry 270(41):24451–24458.

41. Singleton MR, Dillingham MS, Gaudier M, Kowalczykowski SC, Wigley DB (2004) Crystal structure of RecBCD enzyme reveals a machine for processing DNA breaks. Nature 432(7014):187–193.

42. Dillingham MS, Spies M, Kowalczykowski SC (2003) RecBCD enzyme is a bipolar DNA helicase. Nature 423(6942):893–897.

43. Taylor AF, Smith GR (2003) RecBCD enzyme is a DNA helicase with fast and slow motors of opposite polarity. Nature 423(6942):889–893.

44. Emmerson PT (1968) Recombination deficient mutants of Escherichia coli K12 that map between thy A and argA. Genetics 60(1):19–30.

45. Biek DP, Cohen SN (1986) Identification and characterization of recD, a gene affecting plasmid maintenance and recombination in Escherichia coli. Journal of bacteriology 167(2):594–603.

46. Amundsen SK, Taylor AF, Chaudhury AM, Smith GR (1986) recD: the gene for an essential third subunit of exonuclease V. Proceedings of the National Academy of Sciences of the United States of America 83(15):5558–5562.

47. Lovett ST, Luisi-DeLuca C, kolodner RD (1988) The genetic dependence of recombination in recD mutants of Escherichia coli. Genetics 120(1):37–45.

48. Palas KM, Kushner SR (1990) Biochemical and physical characterization of exonuclease V from Escherichia coli. Comparison of the catalytic activities of the RecBC and RecBCD enzymes. Journal of Biological Chemistry 265(6):3447–3454.

49. Datsenko KA, Wanner BL (2000) One-step inactivation of chromosomal genes in Escherichia coli K-12 using PCR products. Proceedings of the National Academy of Sciences of the United States of America 97(12):6640–6645.

50. Baba T et al. (2006) Construction of Escherichia coli K-12 in-frame, single-gene knockout mutants: the Keio collection. Molecular systems biology 2(1):2006.0008.

51. Reisch CR, Prather KLJ (2015) The no-SCAR (Scarless Cas9 Assisted Recombineering) system for genome editing in Escherichia coli. Scientific reports 5:15096.

52. Ennis DG, Levine AS, Koch WH, Woodgate R (1995) Analysis of recA mutants with altered SOS functions. Mutation research 336(1):39–48.

53. Kokubo K, Yamada M, Kanke Y, Nohmi T (2005) Roles of replicative and specialized DNA polymerases in frameshift mutagenesis: mutability of Salmonella typhimurium strains lacking one or all of SOS-inducible DNA polymerases to 26 chemicals. DNA Repair (Amst) 4(10):1160–1171.

54. Reuven NB, Arad G, Maor-Shoshani A, Livneh Z (1999) The mutagenesis protein UmuC is a DNA polymerase activated by UmuD′, RecA, and SSB and is specialized for translesion replication. Journal of Biological Chemistry 274(45):31763–31766.

55. Pagès V, Fuchs RP (2018) Inserting Site-Specific DNA Lesions into Whole Genomes. Methods Mol Biol 1672:107–118.

56. Beloin C et al. (2003) Contribution of DNA conformation and topology in right-handed DNA wrapping by the Bacillus subtilis LrpC protein. J Biol Chem 278(7):5333–5342.

57. Pagès V, Fuchs RPP (2002) How DNA lesions are turned into mutations within cells? Oncogene 21(58):8957–8966.

58. Hashimoto Y, Ray Chaudhuri A, Lopes M, Costanzo V (2010) Rad51 protects nascent DNA from Mre11-dependent degradation and promotes continuous DNA synthesis. Nat Struct Mol Biol 17(11):1305–1311.

59. Schlacher K et al. (2011) Double-Strand Break Repair-Independent Role for BRCA2 in Blocking Stalled Replication Fork Degradation by MRE11. Cell 145(4):529–542.

60. Shevelev IV, Hübscher U (2002) The 3′–5′ exonucleases. Nature reviews Molecular cell biology 3(5):364–376.

61. Lovett ST (2011) The DNA Exonucleases of Escherichia coli. EcoSal Plus 4(2)

62. Schlacher K et al. (2005) DNA polymerase V and RecA protein, a minimal mutasome. Molecular Cell 17(4):561–572.

63. Napolitano R, Janel-Bintz R, Wagner J, Fuchs RP (2000) All three SOS-inducible DNA polymerases (Pol II, Pol IV and Pol V) are involved in induced mutagenesis. The EMBO journal 19(22):6259–6265.

64. Korada SKC et al. (2013) Crystal structures of Escherichia coli exonuclease I in complex with single-stranded DNA provide insights into the mechanism of processive digestion. Nucleic Acids Research 41(11):5887–5897.

65. Lu D, Myers AR, George NP, keck JL (2011) Mechanism of Exonuclease I stimulation by the single-stranded DNA-binding protein. Nucleic Acids Research 39(15):6536– 6545.

66. Viswanathan M, Lovett ST (1998) Single-strand DNA-specific exonucleases in Escherichia coli. Roles in repair and mutation avoidance. Genetics 149(1):7–16.

67. Burdett V, Baitinger C, Viswanathan M, Lovett ST, Modrich P (2001) In vivo requirement for RecJ, ExoVII, ExoI, and ExoX in methyl-directed mismatch repair. Proceedings of the National Academy of Sciences of the United States of America 98(12):6765–6770.

68. Thoms B, Borchers I, Wackernagel W (2008) Effects of Single-Strand DNases ExoI, RecJ, ExoVII, and SbcCD on Homologous Recombination of recBCD+ Strains of Escherichia coli and Roles of SbcB15 and XonA2 ExoI Mutant Enzymes. Journal of bacteriology

69. Belguise-Valladier P, Fuchs RP (1995) N-2-aminofluorene and N-2 acetylaminofluorene adducts: the local sequence context of an adduct and its chemical structure determine its replication properties. J Mol Biol 249(5):903–913.

70. Mazon G et al. (2010) Alkyltransferase-like protein (eATL) prevents mismatch repair-mediated toxicity induced by O6-alkylguanine adducts in Escherichia coli. Proceedings of the National Academy of Sciences 107(42):18050–18055.

71. Bianco PR, Kowalczykowski SC (2000) Translocation step size and mechanism of the RecBC DNA helicase. Nature 405(6784):368–372.

72. Wu CG, Bradford C, Lohman TM (2010) Escherichia coli RecBC helicase has two translocase activities controlled by a single ATPase motor. Nat Struct Mol Biol 17(10):1210–1217.

73. Korangy F, Julin DA (1993) Kinetics and processivity of ATP hydrolysis and DNA unwinding by the RecBC enzyme from Escherichia coli. Biochemistry 32(18):4873– 4880.

74. Curth U, Genschel J, Urbanke C, Greipel J (1996) In vitro and in vivo function of the C-terminus of Escherichia coli single-stranded DNA binding protein. Nucleic Acids Res 24(14):2706–2711.

75. Shereda RD, Kozlov AG, Lohman TM, Cox MM, Keck JL (2008) SSB as an organizer/mobilizer of genome maintenance complexes. Critical reviews in biochemistry and molecular biology 43(5):289–318.

76. Costes A, Lecointe F, McGovern S, Quevillon-Cheruel S, polard P (2010) The C-terminal domain of the bacterial SSB protein acts as a DNA maintenance hub at active chromosome replication forks. PLoS Genetics 6(12):e1001238.

77. Bianco PR (2017) The tale of SSB. Progress in biophysics and molecular biology 127:111–118.

78. Antony E, Lohman TM (2019) Dynamics of E. coli single stranded DNA binding (SSB) protein-DNA complexes. Semin Cell Dev Biol 86:102–111.

79. Bianco PR (2021) The mechanism of action of the SSB interactome reveals it is the first OB-fold family of genome guardians in prokaryotes. Protein Sci 30(9):1757–1775.

80. Shereda RD, Bernstein DA, Keck JL (2007) A central role for SSB in Escherichia coli RecQ DNA helicase function. Journal of Biological Chemistry 282(26):19247–19258.

81. Bianco PR et al. (2017) The IDL of E. coli SSB links ssDNA and protein binding by mediating protein-protein interactions. Protein Sci 26(2):227–241.

82. Thrall ES, Piatt SC, Chang S, Loparo JJ (2022) Replication stalling activates SSB for recruitment of DNA damage tolerance factors. Proc Natl Acad Sci U S A 119(41):e2208875119.

83. Pasero P, Vindigni A (2017) Nucleases Acting at Stalled Forks: How to Reboot the Replication Program with a Few Shortcuts. Annual review of genetics 51:477–499.

84. Patel DR, Weiss RS (2018) A tough row to hoe: when replication forks encounter DNA damage. Biochemical Society transactions

85. Costanzo V (2011) Brca2, Rad51 and Mre11: Performing balancing acts on replication forks. DNA Repair 10(10):1060–1065.

86. Kolinjivadi AM et al. (2017) Moonlighting at replication forks: a new life for homologous recombination proteins BRCA1, BRCA2 and RAD51. FEBS letters

