## supplementary figures for "The RecBC complex protects single-stranded DNA gaps during lesion bypass"

### Co-last authors

*To whom correspondence should be addressed:

**This PDF file includes:**

Table S1

Figures S1 to S7

Supporting Material&Methods

| Name | Short name | Genotype |
|---|---|---|
| EVP22/23 | Parental strain | FBG151/152 *uvrA*::frt *mutS*::frt |
| EVP164/165 | *recB* | FBG151/152 *uvrA*::frt *mutS*::frt *recB::*frt |
| EVP736/737 | *recC* | FBG151/152 *uvrA*::frt *mutS*::frt *recC* ::frt |
| EVP582/583 | *recD* | FBG151/152 *uvrA*::frt *mutS*::frt *recD::*frt |
| EVP669/670 | *exoI (sbcB)* | FBG151/152 *uvrA*::frt *mutS*::frt *exoI::*frt |
| EVP681/682 | *exoI recB* | FBG151/152 *uvrA*::frt *mutS*::frt *exoI*::frt *recB*::frt |
| EVP798/799 | *sbcCD recB* | FBG151/152 *uvrA*::frt *mutS*::frt *sbcCD*::frt *recB::frt* |
| EVP683/684 | *exoVII (xseA) recB* | FBG151/152 *uvrA*::frt *mutS*::frt *exoVII*::frt *recB*::frt |
| EVP685/686 | *exoX recB* | FBG151/152 *uvrA*::frt *mutS*::frt *exoX*::frt *recB*::frt |
| EVP876/877 | *exoIA183V* recB* | FBG151/152 *uvrA*::frt *mutS*::frt *exoIA183V recB*::frt |
| EVP629/630 | *recB1080#* | FBG151/152 *uvrA*::frt *mutS*::frt *recB*D1080A |
| EVP112/113 | *lexA* | FBG151/152 *uvrA*::frt *mutS*::frt *sulA*:: frt *lexA::*frt |
| EVP661/662 | *lexA recB* | FBG151/152 *uvrA*::frt *mutS*::frt *sulA*:: frt *lexA::*frt *recB* ::kan |
| EVP761/762 | *uvrD recB* | FBG151/152 *uvrA*::frt *mutS*::frt *uvrD*::frt *recB*::frt |
| EVP1056/1057 | *dinG recB* | FBG151/152 *uvrA*::frt *mutS*::frt *dinG*::frt *recB*::frt |
| EVP786/780 | *yoaA recB* | FBG151/152 *uvrA*::frt *mutS*::frt *yoaA*::frt *recB*::frt |
| EVP779/781 | *helD recB* | FBG151/152 *uvrA*::frt *mutS*::frt *helD*::frt *recB*::frt |

**Table S1: *E. coli* strains used in this study**

**FBG151 =** MG1655 ΔlacIZ ΔattBλ::aadA-attRλ-'lacZ (lesion inserted in the lagging orientation)
**FBG152 =** MG1655 ΔlacIZ ΔattBλ::aadA-attRλ-'lacZ (lesion inserted in the leading orientation)
***** This mutants has been created by CRISPR-Cas9 technologies, following the protocol of Reisch 2015 {Reisch and Prather, 2015, #623}.
**#** Obtained as explained in {Laureti et al., 2017, #807}

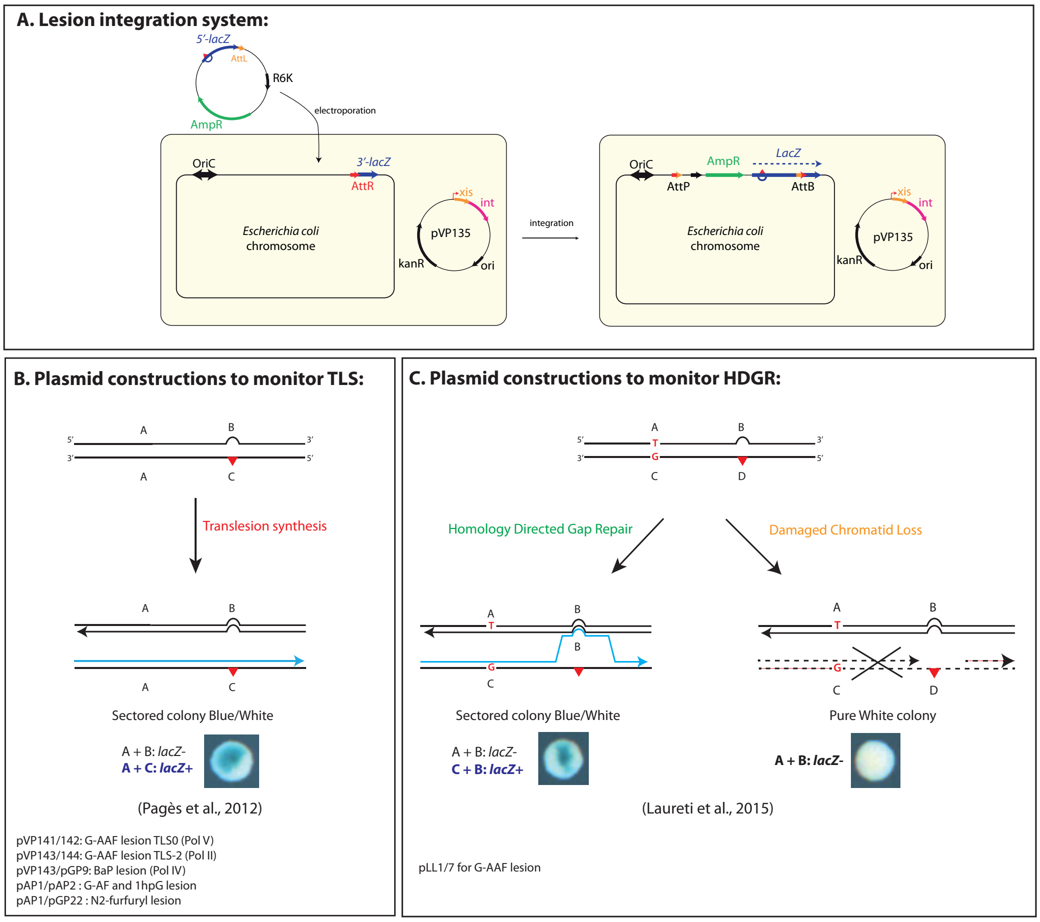


**Fig. S1. Lesion tolerance assay and lesion containing plasmids.**

**A)** The system is based on the phage lambda site-specific recombination and it requires a recipient *E. coli* strain with a single *att*R site fused to the 3'-end of the *lacZ* gene and a non-replicating plasmid construct containing a single DNA lesion (red triangle) in the 5'-end of the *lacZ* gene fused to the *att*L site. The recombination reaction between *att*L and *att*R is controlled by ectopic expression of phage lambda *int*–*xis* proteins (provided by pVP135), and leads to the integration of the lesion-containing vector into the chromosome. Integrants are selected on the basis of their resistance to ampicillin and the chromosomal integration restores an entire *lacZ* gene.

**B)** Plasmid duplexes used to monitor TLS events: the non-damaged strand (A+B markers) contains a short sequence of heterology opposite the lesion that inactivates the *lacZ* gene and serves as a genetic marker for strand discrimination. Only the replication by TLS polymerases of the A+C markers will give rise to a functional *lacZ* gene and therefore to a sectored blue-white colony.

**C)** Plasmid duplexes used to specifically monitor strand exchange mechanisms, *i.e.* HDGR events: four genetic markers have been designed in order to distinguish the replication of the non-damaged strand (containing markers A and B) from the damaged strand (containing markers C and D). Using a combination of frameshift and stop codon, we inactivated *lacZ* gene on both the damaged (C-D) and undamaged (A-B) strands of the vector. Only a strand exchange mechanism by which replication has been initiated on the damaged strand (incorporation of marker C), and where a template switch occurred at the lesion site (leading to incorporation of marker B) will restore a functional *lacZ* gene leading to sectored blue-white colonies. When the damaged strand is lost, only the A and B marker will be replicating giving rise to a white colony.

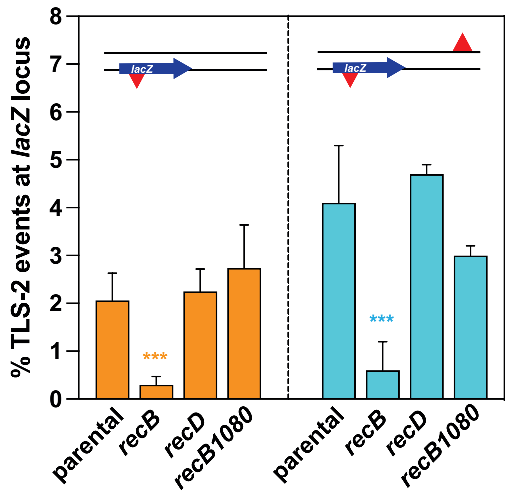


**Fig. S2.** The graph represents Pol II TLS events at a single G-AAF lesion located in the *lacZ* gene in the absence (orange colour) or in the presence of an additional lesion (light blue colour) on the opposite strand. The data in every graph represent the average and standard deviation of at least three independent experiments.

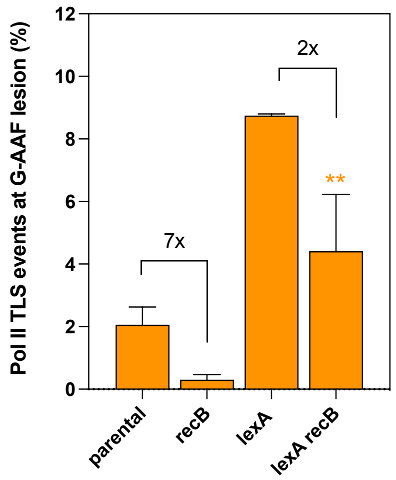


**Fig. S3.** The graph represents the Pol II TLS events in the presence of a G-AAF lesion for the *lexA* and *lexA recB* deficient strain. The data represent the average and standard deviation of more than four independent experiments obtained in the lagging or in the leading orientation. The asterisk represents the statistical significance for the *lexA recB* strain compared to *recB* deficient strain for both TLS events. Statistical analyses have been performed with Prism using an unpaired *t*-test: ** $p < 0,005$.

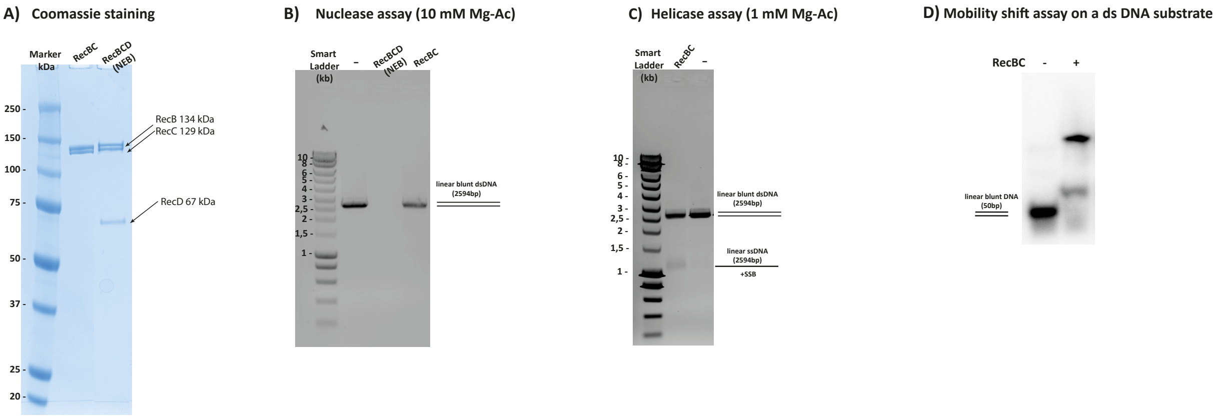


**Fig. S4. A)** Protein gel Coomassie stained showing the purified RecB and RecC proteins in comparison to the RecBCD complex from NEB. **B)** Nuclease assay using a linear blunt dsDNA substrate comparing the activity of RecBC complex versus the RecBCD complex (Provided by NEB), in the presence of 10mM Mg-Ac. **C)** Helicase assay for RecBC complex using a linear blunt dsDNA substrate, in the presence of 1mM Mg-Ac. The difference here from the nuclease assay is the concentration of Mg and the temperature of the reaction (for more details, see Supplementary M&M section). **D)** Mobility shift assay on native acrylamide gel using a linear blunt dsDNA substrate (50 bp), in the presence of 1mM Mg-Ac.

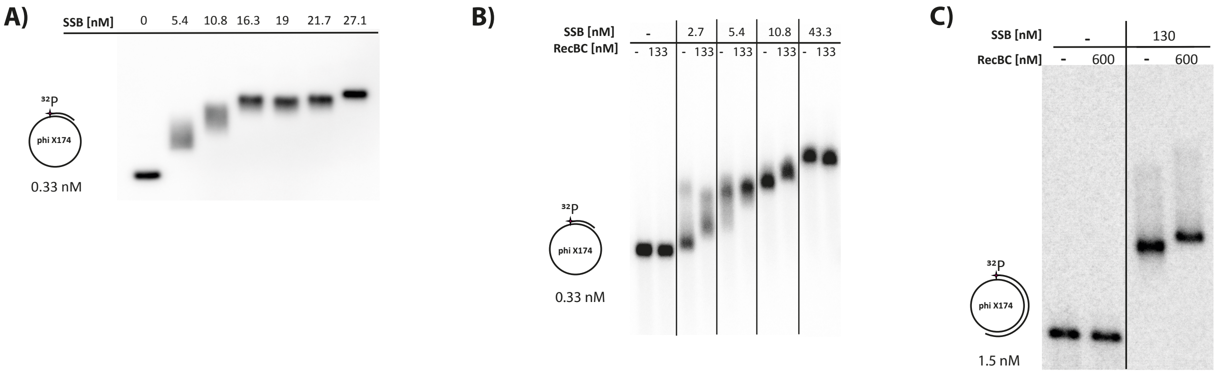


**Fig. S5. A)** Mobility shift assay on agarose gel using the 5.3 kb gapped substrate in the presence of increasing concentrations of SSB. **B)** Mobility shift assay on agarose gel using the 5.3 kb gapped substrate in the presence of a fixed concentration of RecBC complex and with (or without) increasing concentrations of SSB. In the absence of SSB, no band shift is observed for RecBC alone. **C)** Mobility shift assay on agarose gel using the 2.5 kb gapped substrate in the presence of a fixed concentration of RecBC complex and SSB.

**A) ExoI nuclease assay on a gapped substrate**

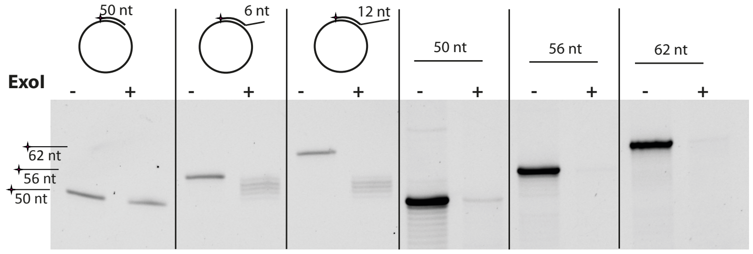


**B) Helicase genetic screen**

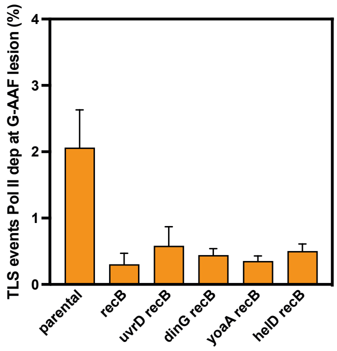


**C) ExoI nuclease assay on a oligo**

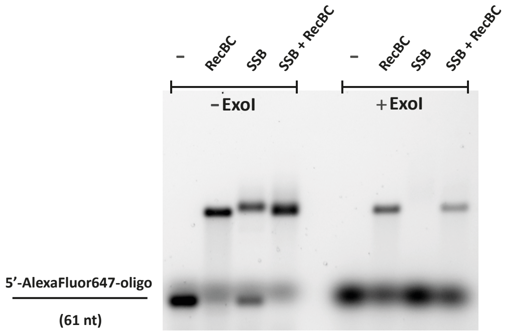


**Fig. S6. A)** On the left of the denaturing 15% acrylamide gel: ExoI nuclease assay using a gapped substrate without flap (50 nt oligo), with a 6nt flap and with a 12 nt flap, respectively. On the right of the gel, ExoI nuclease assay in the presence of three different oligos (50, 56, 62 nt) as control. **B)** The graph represents the Pol II TLS events in the presence of a G-AAF lesion for several helicase genes in the *recB* deficient strain. The data represent the average and standard deviation of more than four independent experiments obtained in the lagging or in the leading orientation. No statistically significant difference was observed. **C)** On the left mobility shift assay using a 61nt fluorescent oligo in the presence of RecBC, SSB and RecBC+SSB, respectively. On the right side the complexes formed were digested by ExoI. In the presence of RecBC the oligo is protected.

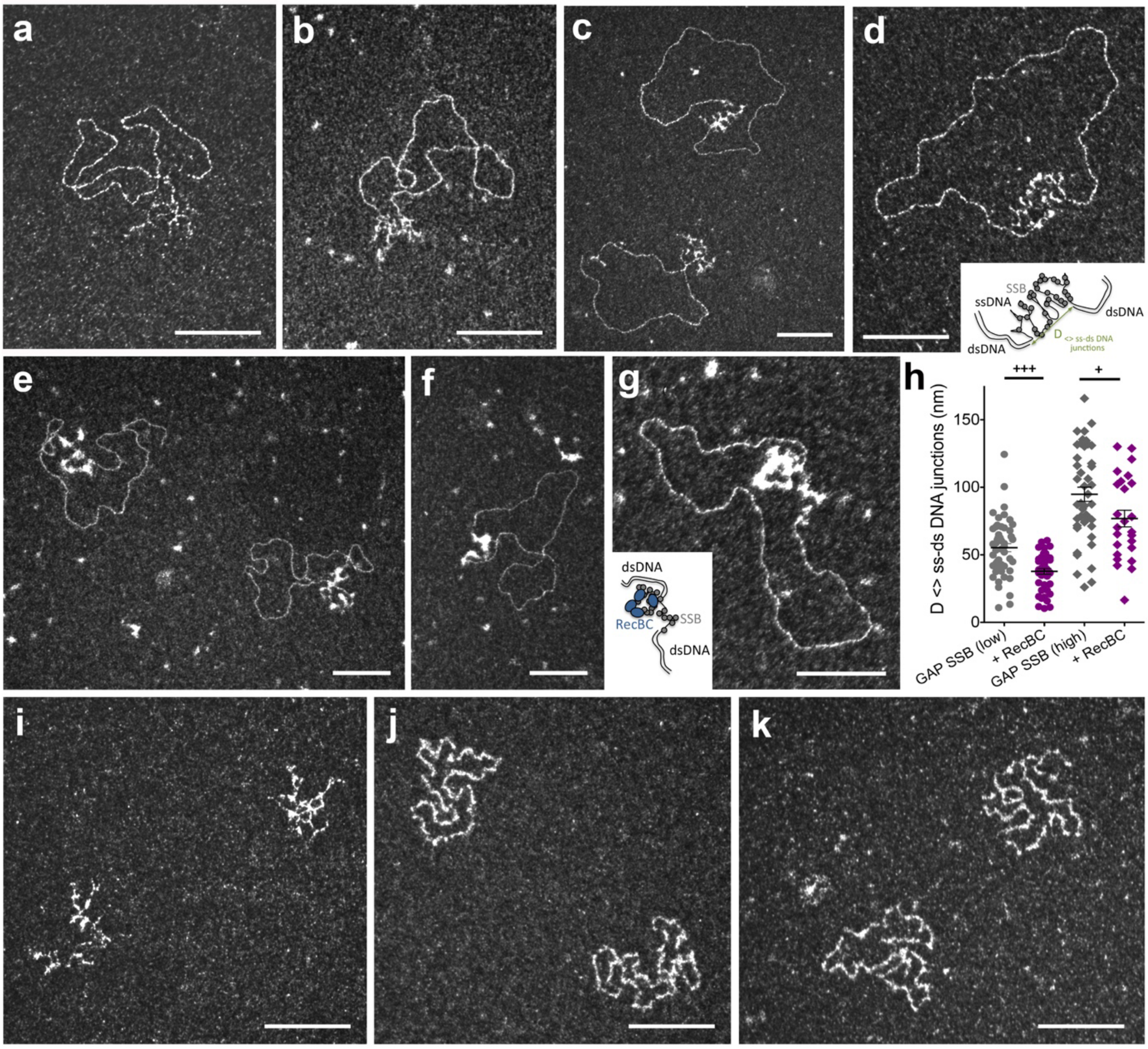


**Suppl Figure 7. Transmission Electron Microscopy analysis of RecBC-SSB-ssDNA complexes in darkfield imaging mode. a)** Control DNA substrate containing a 2.5 kb ssDNA gap. ssDNA is folded into secondary structures under these spreading conditions. **b**) 300 nM RecBC were incubated with the 2.5 kb ssDNA gapped substrate (10 μM nucleotides - 3.2 nM in molecules). RecBC does not bind to the substrate in absence of SSB. **c-d)** A low SSB concentration (135 nM) was incubated with the 2.5 kb ssDNA gapped substrate (10 μM nucleotides - 3.2 nM in molecules). The ssDNA part of the substrate is partially covered by SSB protein. **e-g)** 200 nM of RecBC were added to the reaction. RecBC binding to SSB-ssDNA induces its compaction. **h)** Distribution of the distance between the two ss-dsDNA junctions showing that in both SSB concentrations (135 and 286 nM), RecBC reduces then compacts SSB-ssDNA length. The statistical unpaired t-test was applied, *** corresponded to a Pvalue $P < 0.001$. **i)** TEM representative view of the Phix 174 ssDNA virion. **j)** 10 μM of the PhiX 174 virion is incubated with 286 nM SSB. SSB partially covers the ssDNA. **k)** 200 nM RecBC are added to the SSB-ssDNA shown in j). RecBC does not bind to SSB-ssDNA but appears in the background. All scale bars correspond to 100 nm.

**SUPPLEMENTARY MATERIAL AND METHODS**

**RecBC nuclease assay**

5.8 nM of a linearized plasmid (2594 bp) was incubated 30’ at 37°C in 50 mM K-Ac, 20 mM Tris-Ac, 10 mM Mg-Ac, 1 mM DTT, 1 mM ATP with 30 nM RecBC or RecBCD (ratio protein:DNA 5:1). The samples were loaded on 1% agarose gel with ethidium bromide and visualized with Chemidoc MP (Biorad). RecBCD was provided by NEB.

**RecBC helicase assay**

5.3 nM of a linearized plasmid (2594 bp) was incubated 30’ at 25°C in 50 mM K-Ac, 20 mM Tris-Ac, 1 mM Mg-Ac, 1 mM DTT, 1mM ATP, 0.36 µM SSB with 27 nM RecBC (ratio protein:DNA 5:1). The samples were loaded on 1% agarose gel with ethidium bromide and visualized with Chemidoc MP (Biorad).

**Electrophoretic mobility shift assay on a dsDNA substrate**

2 nM of a $^{32}P$ 5’-radiolabelled 50 bp dsDNA was incubated 20’ at 25°C in 10 µl binding buffer (20 mM Tris-HCl pH8, 50 mM NaCl, 100 mg/ml BSA-Ac, 1 mM DTT and 1 mM Mg-Ac) with 600 nM RecBC. The samples were loaded on 12% native acrylamide gel. The gel was dried and scanned with Typhoon (Cytiva).

**Electrophoretic mobility shift assays on a gapped substrate**

For the experiment in Suppl Fig 5A, 0.33 nM of the radiolabelled 5.3 kb gapped substrate (1.7 µM nucleotides for the ssDNA) were incubated for 20’ at RT with increasing concentrations of SSB tetramer (0 – 5.4 – 10.8 – 16.3 - 19 -21.7 – 27.1 nM) in 12 µl binding buffer. The samples were loaded on 0.8% agarose gel. The gel was dried for 1h30 at 60°C, exposed overnight to a Fuji screen and scanned with a Typhoon (Cytiva).

For the experiment in Suppl Fig 5B, 0.33 nM of the radiolabelled 5.3 kb gapped substrate were first incubated for 20’ at RT with with increasing concentrations of SSB tetramer (0 – 2.7 – 5.4 – 10.8 – 43.3 nM) in 12 µl binding buffer. Then 133 nM RecBC (or RecBC buffer as a negative control) was added to the reaction and incubated for 20’ at RT. The samples were loaded on 0.6% agarose gel, running at 4°C. The gel was dried for 1h30 at 60°C, exposed overnight to a Fuji screen and scanned with a Typhoon (Cytiva).

For the experiment in Suppl Fig 5C, 1.5 nM of the radiolabelled 2.5 kb gapped substrate (3.7 µM nucleotides for ssDNA) were first incubated for 20' at RT with 0 or 130 nM of SSB tetramer in 10 µl binding buffer. Then 600 nM RecBC (or RecBC buffer as a negative control) was added to the reaction and incubated for 20' at RT. The samples were loaded on 0.6% agarose gel, running at 4°C. The gel was dried for 1h30 at 60°C, exposed overnight to a Fuji screen and scanned with a Typhoon (Cytiva).

**Electrophoretic mobility shift assay and ExoI nuclease assay on an oligonucleotide**

10 nM of a 61-mer oligonucleotide 5'-labelled with Alexa Fluor 647 was incubated 20' at 25°C in 10 µl binding buffer with 5.5 nM SSB or 160 nM RecBC or both proteins. Half of each reaction was digested by 45 nM ExoI (15', 37°C). The samples were loaded on 1.5% agarose gel and visualized with Chemidoc MP (Biorad).

**ExoI nuclease assay**

3 different oligonucleotides 5'-labelled with FAM (VP770, VP1332 and VP1334) were hybridized to φX174 ssDNA plasmids to create gapped-plasmids without flap, with a 6 nt flap and with a 12 nt flap, respectively. 6 nM of each substrate was digested by 34 nM ExoI (NEB) during 30' at 37°C. Reactions were stopped by addition of formamide dye. The samples were loaded on denaturing 15% acrylamide gel and visualized with Chemidoc MP.
